# A change in behavioral state switches the pattern of motor output that underlies rhythmic head and orofacial movements

**DOI:** 10.1101/2022.12.29.522264

**Authors:** Song-Mao Liao, David Kleinfeld

## Abstract

The breathing rhythm serves as a reference that paces orofacial motor actions and orchestrates active sensing. Past work reports that pacing occurs solely at a fixed phase relative to sniffing. We reëvaluated this constraint as a function of exploratory behavior. Allocentric and egocentric rotations of the head and the electromyogenic activity of the underlying motoneurons for head and orofacial movements were recorded in free-ranging rats as they searched for food. We found that a change in state from foraging to rearing is accompanied by a change in the phase of muscular activation relative to sniffing, so that pacing now occurs at one of two phases. Further, head-turning is biased such that an animal gathers a novel sample of its environment upon inhalation. In toto, the coordination of active sensing has a previously unrealized computational complexity that, in principle, can emerge from hindbrain circuits with fixed architecture and credible synaptic time-delays.

## INTRODUCTION

Many natural behaviors in animals involve significant rhythmic components. First and foremost, these encompass locomotion and orienting, the essential motor actions for navigation, through the coordination of brainstem and spinal motor plants.^1-3^ The kinematics of locomotion are well studied^4^, and the dynamics are computationally rich. In particular, the relative timing among the muscle groups that drive the limbs in tetrapeds will change as the speed of the animal increases and the gait progresses from walking to jogging to trotting to cantering and lastly to galloping.^5,6^ This can be solely enabled by phase-shifts that accumulate in networks of oscillators with time-delayed connections.^7-11^ A similar dynamic occurs in networks that govern undulatory locomotion.^12^ In contrast to the case of locomotion, relatively little is known about the kinematics and dynamics of orienting motor actions. One claim that motivates the present work is that vertical head bobbing, i.e., changes in pitch, is phase-locked to breathing as rodents explore their environment.^13-15^

Beyond head-bobbing, the breathing rhythm plays an outsized role in the coordination of orofacial motor actions in rodents.^13,16^ Quantitative measurements of the coordination of orofacial motor control by breathing were reported for whisking^17-22^, facial movement via the mystacial pad^17^, and nose twitching^14^ with freely moving rodents that inhabited a small platform. In all cases, the movement was locked to a single phase in the breathing cycle. These results led to the “master oscillator” hypothesis, where breathing serves as the reference oscillator to bind orofacial sensory inputs.^16,23,24^ In principle, this permits sensory information from multiple modalities to be gathered with a precise temporal relationship for efficient and accurate neuronal computations. In fact, recent evidence^15,25^ supports the role of the respiration cycle as a reference oscillator in high-level sensory and cognitive processing.^26,27^ Yet, all known past studies on the phase of motor actions with respect to breathing, and on the phase of computational actions with respect to breathing, report that locking occurs solely at a single, fixed phase.^13,14,17,28,29^ This implies a surprisingly harsh constraint on the combinatorics that govern the coordination of different actions into behaviors.

We believe that the sparsity of ethological context in the foundational studies^13-22,25-27^ had the untold effect of limiting our understanding of a richer computational complexity. We thus hypothesize that the coordination of motor actions with breathing can exhibit multiple, stable phase relations when rodents can switch among different ethological contexts. We test this hypothesis using a behavioral task that involves exploration, navigation, and active sensation as rats search for and ingest food in a large arena. We ask: (*i*) Does the frequency of sniffing change with behavioral state? This question is motivated by a change in sniffing frequency between odor sampling and reward selection in a forced-choice task.^30^ (*ii*) Does the coordinated constriction among neck and orofacial muscles relative to themselves, as well as to breathing, depend on behavioral state? This question is motivated by changes in gait concurrent with changes in the speed of locomotion.^31,32^ (*iii*) How does activation of the musculature involved in head-turning and nose turning coordinate with breathing, as well as respect the midline as an axis of symmetry? This addresses the question of when should rodents best sample their environment as they rotate their head and nose outward with respect to their midline. Motivated by prior work on horizontal head movement in mice^33^, we monitored head orientation and head-turns relative to the torso, along with the activation of muscles that drive head motion and the orofacial actions of the nose and face.

## RESULTS

We trained 33 food-restricted rats to search in the dark for food within an open circular arena of 1 m in diameter (**Figure 1A**). A 9-axis absolute orientation sensor is mounted onto the skull to measure acceleration of the head in Euclidian coordinates, the 3-dimensional angular velocity in Euler coordinates, and the absolute orientation, i.e., yaw, pitch, and roll, at 100 Hz (**Figure 1B**); these define movement in allocentric coordinates. In 17 animals, a second absolute orientation sensor was embedded subcutaneously to measure movement and orientation of the torso (**Figure 1B**). The torso sensor signals are subtracted from the head sensor signals to obtain the relative head-torso orientation and angular velocities; these define movement in egocentric coordinates. The difference in sensor orientation was found to drift by less than 0.01° over the period of recording. In all animals, respiration was recorded by an implanted thermocouple in the nasal cavity that measured the cyclic change in air temperature concurrent with inhalation and expiration (**Figure 1B**). We used a Hilbert transform to locate the onsets of inspiration, which we took as the start of each breathing cycle with a phase of 0 radians. Videography was used to track the position of the animal in the arena (**Figure 1B,C**).

**Figure 1.**
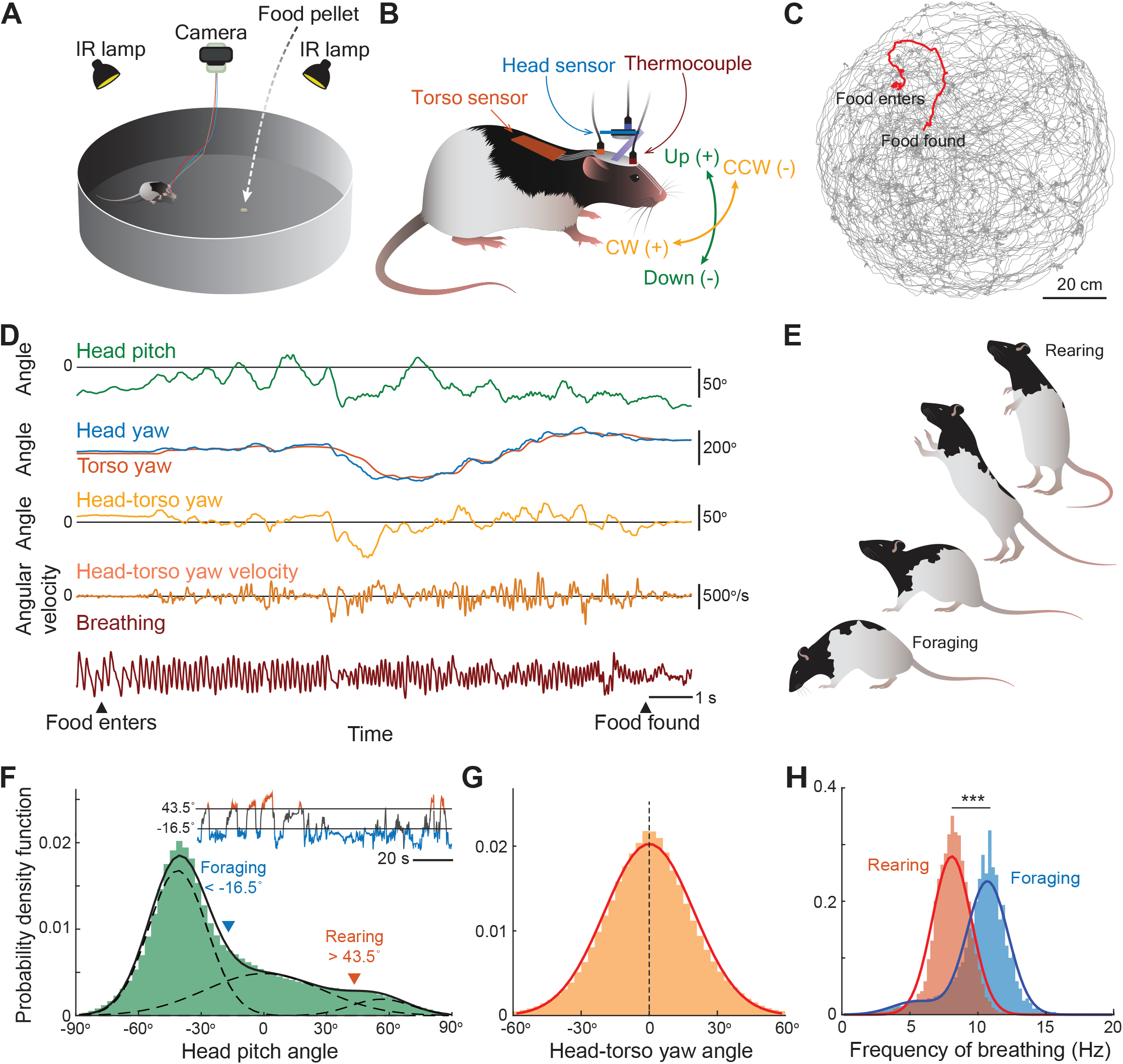
Head movements and breathing in foraging and rearing rats. **(A)** Depiction of the foraging task in the circular arena, 1 meter in diameter. **(B)** Depiction of movements of the head and torso that are measured with orientation sensors and breathing with a thermocouple from freely moving rats. We define the horizontal (yaw) angular velocity to be positive when the rotation is clockwise (CW) and negative when the rotation is counterclockwise (CCW), as viewed from above. The vertical (pitch) angle is defined as positive when the head is raised up against the gravity and is defined as negative when the head is tilted downward. **(C)** Trajectories of 72 foraging trials for one rat as traced by DeepLabCut. The path in panel D is highlighted in red. **(D)** Example data of the pitch angle of the head, the yaw angle of the head, torso, and head-torso, the yaw velocity of head-torso, and breathing in one foraging rat. Left arrowhead indicates the time when the food pellet is dropped into the arena and the right arrowhead the time when the rat finds the food. We choose the polarity of the thermocouple signal so that increasing values, i.e., rising, positive slope, indicates inhaling and decreasing values, i.e., negative slope, indicates exhaling. **(E)** Illustration of rodent behaviors that are formed with foraging and rearing states. **(F)** Probability density distribution of the head pitch angle. Data pooled from 29 rats. Distribution is fitted by three Gaussians. Intersections of the Gaussian fits at -16.5° and 43.5° are labelled and used as the thresholds to define the behavioral states. Insert: Example time series of the pitch angle; see also **FIgure S1A**. **(G)** Probability density distribution of the head-torso yaw angle. Data pooled from 17 rats. Red line indicates the Gaussian fit (mean = 0, SD = 19.7°). **(H)** Probability density distribution of the periods of breathing cycles in foraging and rearing states. Data pooled from 33 rats. Rearing and foraging distributions are fitted with one and two Gaussians respectively. Mean, standard deviation, and relative amplitude of the fit (mean, SD, relative amplitude) are: (8.1 Hz, 1.4 Hz, 1.0) for rearing; (5.5 Hz, 1.8 Hz, 0.10) and (10.7 Hz, 1.5 Hz, 0.90) for foraging.

### Behavioral states are revealed when animals search inside an arena

Small, regularly-sized pellets of food were individually dropped at a random location in the arena after an auditory clue (**Figure 1A**). The rats were observed to travel along approximate curvilinear arcs that spanned the entire arena as they searched for a pellet and then lapped the pellet with their tongue and swallowed. After a brief delay, the process started anew. Each rat executed rich head movements in both the horizontal (yaw) and vertical (pitch) directions in the search for food. **Figure 1D** shows an example measurement of head-torso movement and breathing in a trial where the animal was locomoting and sniffing with its head downward to the floor in an act designated as foraging (**Figure 1E**). On occasion, the rat paused its locomotion and rose up on its hind legs to head-bob and sniff in an act designated rearing (**Figure 1E**). We used the instantaneous head pitch angle to categorize these behavioral states across a population (29 rats) in terms of the probability density of the head-pitch angle (**Figure 1F**). The distribution is well fit with three Gaussians (r^2^ = 0.983; **Figure 1F**). The intersections between neighboring Gaussians provides a conservative metric to define a threshold pitch for the foraging state, i.e., head pitch below –16.5°, and a threshold for the rearing state, i.e., head pitch above 43.5° (**Figures 1F** and **S1A**). Lastly, while the distribution of head pitch angles is multimodal (**Figure 1F**), the distribution of the head-torso yaw angle, i.e., head-turning in the horizontal plane, is well fit by a single Gaussian distribution that is centered at zero (r^2^ = 0.992; **Figure 1G**).

ö The distribution of sniffing frequencies associated with rhythmic head-torso turning was found to depend on the behavioral state of the rat (**Figure 1H**). Foraging occurs with a frequency centered near 11 Hz, while rearing occurs at a lower frequency centered near 8 Hz. The separation of center frequencies for the two states is significant (p < 0.001 across 32 of 33 rats). All told, foraging is defined by a low pitch-angle and a higher breathing rate, while rearing is defined by a high pitch-angle and a lower breathing rate (**Figure 1E,F,H**).

### Head movements phase-locked to breathing during both foraging and rearing

Past work established that changes in the pitch of an animal’s head while it explores is locked to the breathing rhythm.^14^ Here we further observed that head-torso turning in the horizontal plane, i.e., egocentric movement, is rhythmic with two spectral components. The movement involves the waxing and waning of a slow spectral component below 5 Hz, the frequency band for orienting and basal breathing, as well as a fast component centered near 10 Hz (**Figure 2A**), the frequency of sniffing (**Figure 1H**). These spectral components also appear in absolute rotations in head-yaw, i.e., allocentric movement, along with head-pitch and breathing (**Figure 2B**). To quantify the relation of egocentric yaw relative to breathing, we calculated the spectral coherence between head-torso yaw and breathing, a measure of how well these two rhythms track over time. Indeed, the two rhythms are strongly phase-locked in the 8 - 12 Hz band, albeit weakly phase-locked in the 1 - 5 Hz band, that encompasses rhythmic head-turning. Here and everywhere phase-locking is defined as statistically significant coherence (**Figure 2A,C**).

**Figure 2.**
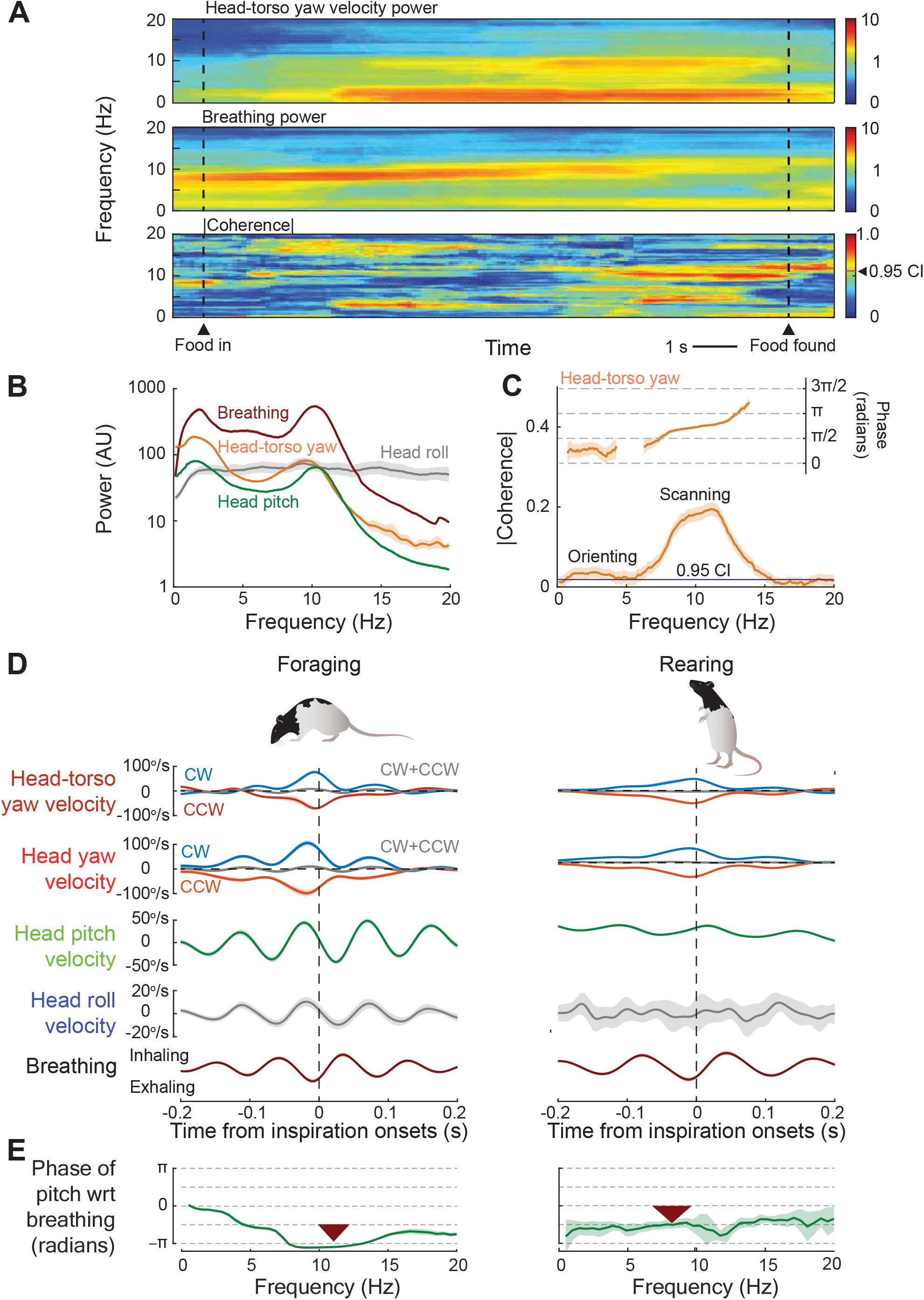
Head movement is phase-locked to breathing with a phase modulated by posture. **(A)** Spectrogram of the head-torso yaw velocity, i.e., power in the velocity versus time (top), spectrogram of breathing (middle), and their spectral coherence (bottom) from data shown in **Figure 1D**. Red band in the coherence indicate close tracking of yaw velocity and breathing. Spectra and coherence are calculated with a moving window of width 5 s and a 0.1 s step size. The half-bandwidth is 1 Hz (9 tapers) and the 0.95 confidence level is |C| = 0.56. **(B)** Spectra of breathing, head pitch velocity, head roll velocity, and head-torso yaw velocity, shown with jackknife error bars. Data from one rat; 582 windows of size = 4 s, half-bandwidth = 1 Hz (7 tapers). Note peaks at 2 Hz and 11 Hz, the later signal shows that head movement is phase-locked to breathing. **(C)** Coherence of head-torso yaw velocity with breathing during foraging. Data, from the same rat as in panel B, shown with jackknife error bars; 582 windows of 4 s in width, half-bandwidth of 2 Hz (15 tapers). Horizontal line indicates 0.95 confidence level. **(D)** Summary of egocentric head-yaw and allocentric head movements relative to the onset of inspiration. Error bars are standard errors across rats. Data of egocentric movements pooled from 17/15 rats in the foraging/rearing states. Data of allocentric movements pooled from 33/31 rats in the foraging/rearing states. Two rats did not have rearing epochs longer than 0.4 s for averaging. **(E)** Phase of the statistically significant regions of the coherence between head pitch velocity and breathing using the data of panel D; the half-bandwidth is 2 Hz. Note the shift in phase of about π/2 radians between foraging and rearing, which shows that phase is modulated by posture. The arrows point to the center frequency of sniffing, and the width is the FWHM of the associated distribution (**Figure 1H**).

Is the timing between head movements and breathing the same during foraging versus rearing? We considered egocentric head-torso turns and all allosteric head turns, divided the time series of movements into epochs with clockwise (CW) versus counter-clockwise (CCW) yaw movements, and calculated the correlation between the angular velocity of each movement and breathing (**Figure 2D**). We observed that head-torso yaw angular velocity is rhythmic and phase-locked to breathing for both foraging and rearing. The peak amplitude occurs close to that of the onset of inspiration for both CW and CCW movements and the time-lag is greater for foraging than rearing (**Figure 2D**). Lastly, phase-locking is also observed for allocentric head yaw-angular velocity and breathing (**Figure 2D**). As a control, the correlation between yaw movement and breathing is lost when the sum of the two directions is considered (black curve, **Figure 2D**), confirming that the rat makes approximately equal numbers of CW versus CCW movements in the foraging state; (N_CCW_ - N_CW_)/(N_CCW_ + N_CW_) = -0.004 ± 0.018 for egocentric head turning (N_CCW_ and N_CW_ calculated over all trials for each of 17 rats, with 254 trials total, and the ratio is averaged over all rats) and -0.017 ± 0.028 for allocentric head turning (471 total trials and an average across 33 rats).

The timing for the angular velocity of head-pitch relative to breathing advances from roughly a quarter-cycle after the onset of inspiration to roughly a quarter-cycle before the onset. This is half-a cycle in total or equivalently π radians in phase, as the rat changes state from foraging to rearing (**Figure 2E**). In contrast to the cases of yaw and pitch, the correlation of head-roll angular velocity with breathing is apparent during foraging but is essentially absent during rearing (gray curve, **Figure 2D**). All told, these data show that changes in behavioral state lead not only to a shift in the frequency of breathing (**Figure 1H**), but also a shift in the phase relation between breathing and both egocentric and allocentric movement of the head (**Figure 2C-E**).

### Neck muscles phase-locked to breathing during both foraging and rearing

The movement of the head is controlled by a multiplicity of muscles.^34^ To record the motor outputs directly from the musculature that contributes to the sniffing-correlated head movements, we measured the electromyogram (EMG) from three ventrolateral neck muscles, i.e., the sternomastoid (SM) (3 bilaterally and 4 unilaterally implanted rats), cleidomastoid (CM) (4 bilateral and 4 unilateral), and clavotrapezius (CT) (6 bilateral and 5 unilateral), and two dorsal neck muscles, i.e., the splenius (SP) (6 bilateral and 4 unilateral) and biventer cervicis (BC) (3 bilateral and 4 unilateral) (**Figure 3A**). **Figure 3B-D** shows three example sets of raw and processed EMG data, the latter to indicate the envelope of muscle excitation, along with egocentric head movement and breathing. In general, we obtained between 3 and 6 measurements for each of the five muscle groups across the two sides of the body (**Table S1**).

**Figure 3.**
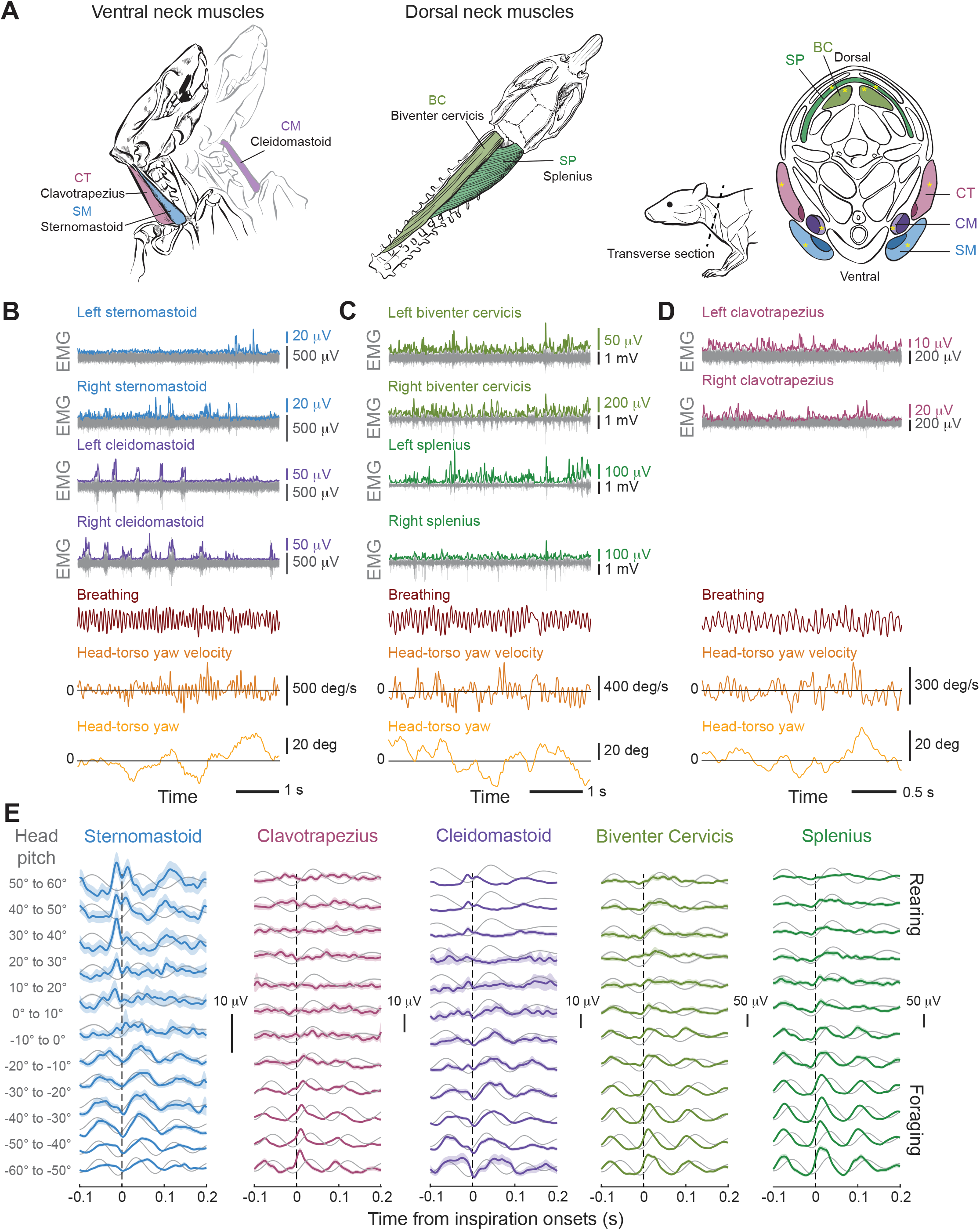
Electromyogenic recording from neck muscles during full arena search. **(A)** (Left) Anatomical depiction of ventral neck muscles sternomastoid (SM), cleidomastoid (CM), and clavotrapezius (CT). (Middle) Anatomical depiction of dorsal neck muscles splenius (SP) and biventer cervicis (BC). (Right) Transverse section (dashed line) showing the relative locations of the muscles. Yellow asterisks indicate the locations where the electromyogram (EMG) signals are recorded. Illustrations adopted from the work of Peterson ^60,61^. **(B)** Example EMGs from bilateral sternomastoid and cleidomastoid muscles (gray), along with the processed data to yield the envelope of muscle activation, and the concurrent breathing and head-torso yaw movements. Data from one rat. **(C)** Example EMGs from bilateral biventer cervicis and splenius muscles (gray); otherwise as above. **(D)** Example EMGs from bilateral clavotrapezius; otherwise as above. **(E)** Cross-correlation of the EMG envelopes of all five neck muscles with respect to the onset of inspiration (sniffing, 4 - 14 Hz) and as a function of head-pitch angle. Shaded areas indicate the standard errors. Data from individual rats. Number of inspiration onsets (foraging/uncategorized/rearing): SM (4,098/1,186/380); CM (2,671/862/1,030); CT (2,632/906/368); SP (4,836/1,122/473); and BC (5,199/808/144). Except for the sternomastoid muscle, correlations are stronger during foraging. The gray traces are the autocorrelations of sniffing.

The activities of all five neck muscles were phase-locked with head movements (**Figure S1B-D**), thus confirming their individual roles as drivers of head movement. We further observed that the activation of all five neck muscles were correlated with the onset of sniffing (**Figure 3E**). The detailed pattern of activity was modulated by the rat’s posture, i.e., its head-pitch angle (**Figure 3E**). The sternomastoid muscle is unique in that it has the greatest amplitude of modulation for rearing, in concordance with the associated change in pitch. All other ventral and dorsal muscles have a greater amplitude and present a simpler rhythmic pattern during foraging as opposed to rearing.

### Vibrissa, facial, and nose muscles lock to breathing for both foraging and rearing

We revisited the coordination of three orofacial motor actions with breathing, i.e., whisking^14,17^, retraction of the mystacial pad^17^, and nose twitching^14,35^, in light of our finding that different patterns of movement and muscular control are revealed by changes in posture (**Figures 2** and **3**). For whisking, we recorded the EMG from the intrinsic protractor (VI) muscles^36-38^ (5 rats) (**Figure 4A**). For movement of the pad, we recorded the EMG from the extrinsic vibrissa muscle nasolabialis^36,39^ (NL) (3 rats) (**Figure 4A**), which further retracts the vibrissae by pulling their exit point from the follicles toward the back.^36^ To measure nose twitching, we recorded the EMG from the deflector nasi (DN) muscle^40^ (7 rats) (**Figure 4A**). This muscle pulls the nose upward and lateral under ipsilateral activation and pulls solely upward under bilateral coactivation.^14^ **Figure 4B,C** shows two sets of example EMG recordings, along with breathing and head movements as the rat foraged. In general, we obtained 3 or 4 measurements for each of these three muscle groups across the two sides of the body (**Table S1**).

**Figure 4.**
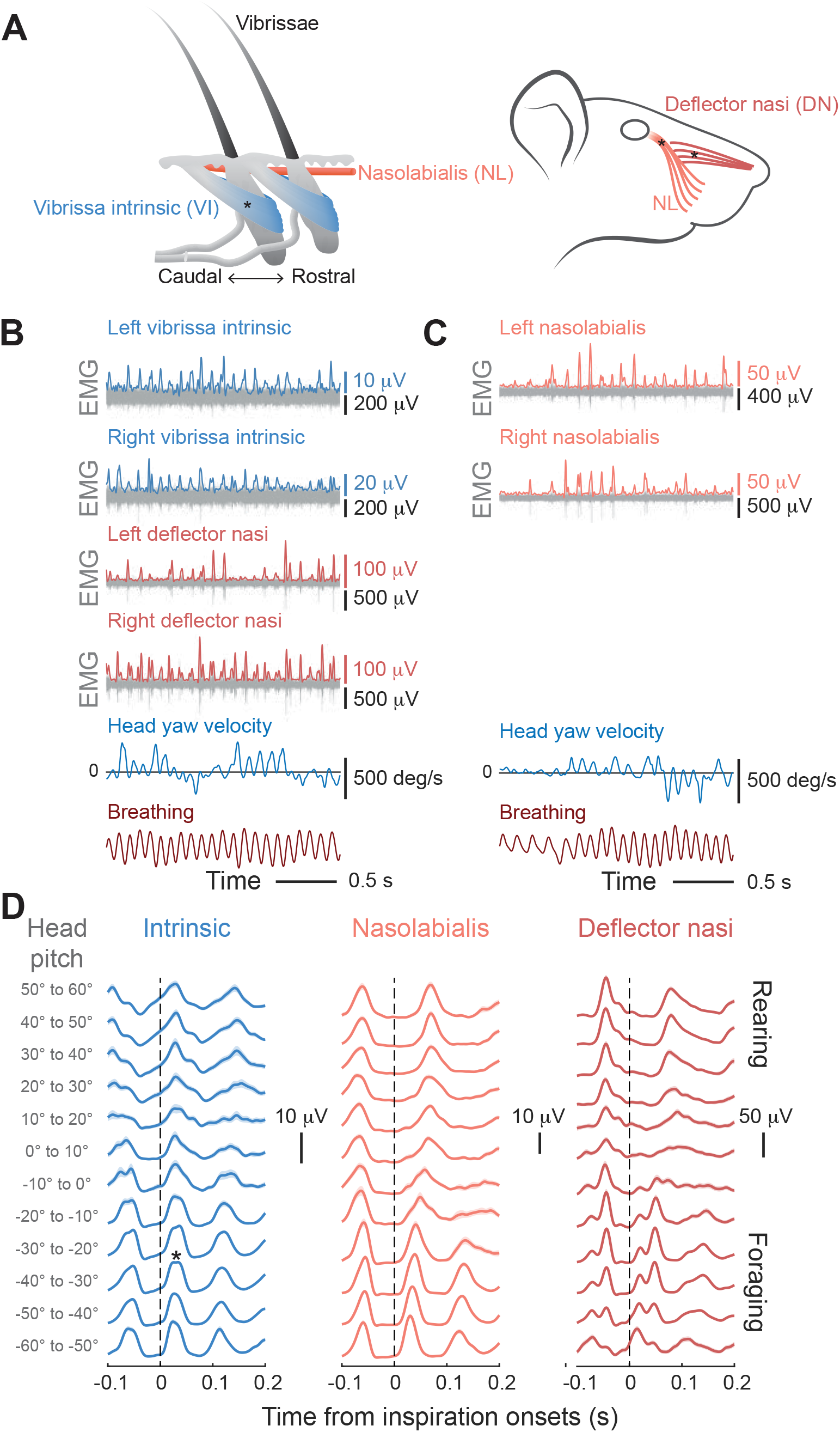
Coordination of orofacial muscles with breathing during full arena search. **(A)** Illustration of the anatomy of vibrissa intrinsic (VI), nasolabialis (NL), and deflector nasi (DN) muscles. Asterisks indicate the approximate locations where the EMG is recorded. **(B)** Example EMGs of bilateral intrinsic and deflector nasi muscles (gray), along with the processed data to yield the envelope of muscle activation, and the concurrent breathing and head-yaw movements. Data from one rat. **(C)** Example EMGs of bilateral nasolabialis muscles (gray); otherwise as above. **(D)** Cross-correlation of the EMG envelopes of VI, NL, and DN muscles with respect to the onset of inspiration (sniffing, 4 – 14 Hz) and as a function of head-pitch angle. Shaded areas indicate the standard errors. Data from individual rats. The timing of the whisking protractor (VI) is essentially unchanged as the rat changes state, while that for nose wiggling (DN) has the greatest change with state. Number of inspiration onsets (foraging/uncategorized/rearing): VI (5,146/1,391/801); NL (4,670/2,303/472); and DN (3,999/1,622/987).

Similar to the neck muscles (**Figure 3E**), we observed a shift in the timing of activation of the orofacial muscles with the onset of inspiration as a function of the pitch-angle (**Figure 4D**). A change in the shape of the waveform from sine- to sawtooth-like is seen for intrinsic muscle activity (3 rats) in the transition from foraging to rearing, i.e., increasing pitch-angle, although timing is largely unchanged. In contrast to the case for the intrinsic muscle, nasolabialis activity (3 rats) and deflector nasi activity (7 rats) are retarded in time across the transition from foraging to rearing. All told, a change in behavioral state from foraging to rearing affects the timing of orofacial (**Figure 4**) as well as orienting (**Figure 3**) motor actions.

### Shifts in phase among motor actions and breathing between foraging and rearing

To gain further insight into the temporal changes in muscular control between the states of foraging versus rearing, we divided the EMG data for the five neck muscles according to the state of animals: the foraging state, with head pitch angle less than – 16.5°, and the rearing state, with angle > 43.5° (**Figure 1F**); responses at the intermediate angles between –16.5° and 43.5° were not further characterized. These data sets were found to cluster into two subgroups. The first subgroup consists of the sternomastoid and cleidomastoid muscles. During foraging, these two muscles are maximally activated upon exhalation (**Figure 5A**). When the animal switched from the foraging to the rearing state, the activities of these muscles dramatically shift and now coincide with the onset of inspiration. The second subgroup consists of the clavotrapezius, splenius, and biventer cervicis muscles. These three muscles are maximally activated just after the onset of inspiration as the rat forages, with a phase difference of about -0.75π radians compared to the first subgroup (**Figure 5A**). Here, when the animal switched from foraging to rearing, the modulation of the clavotrapezius, splenius, and biventer cervicis muscles by breathing is diminished and shifted toward a peak modulation at expiration. The phase difference between muscle subgroups is maintained. All told, these data show that the behavioral state will change the motor output of specific muscles that set the phase of rhythmic head movement with respect to breathing.

**Figure 5.**
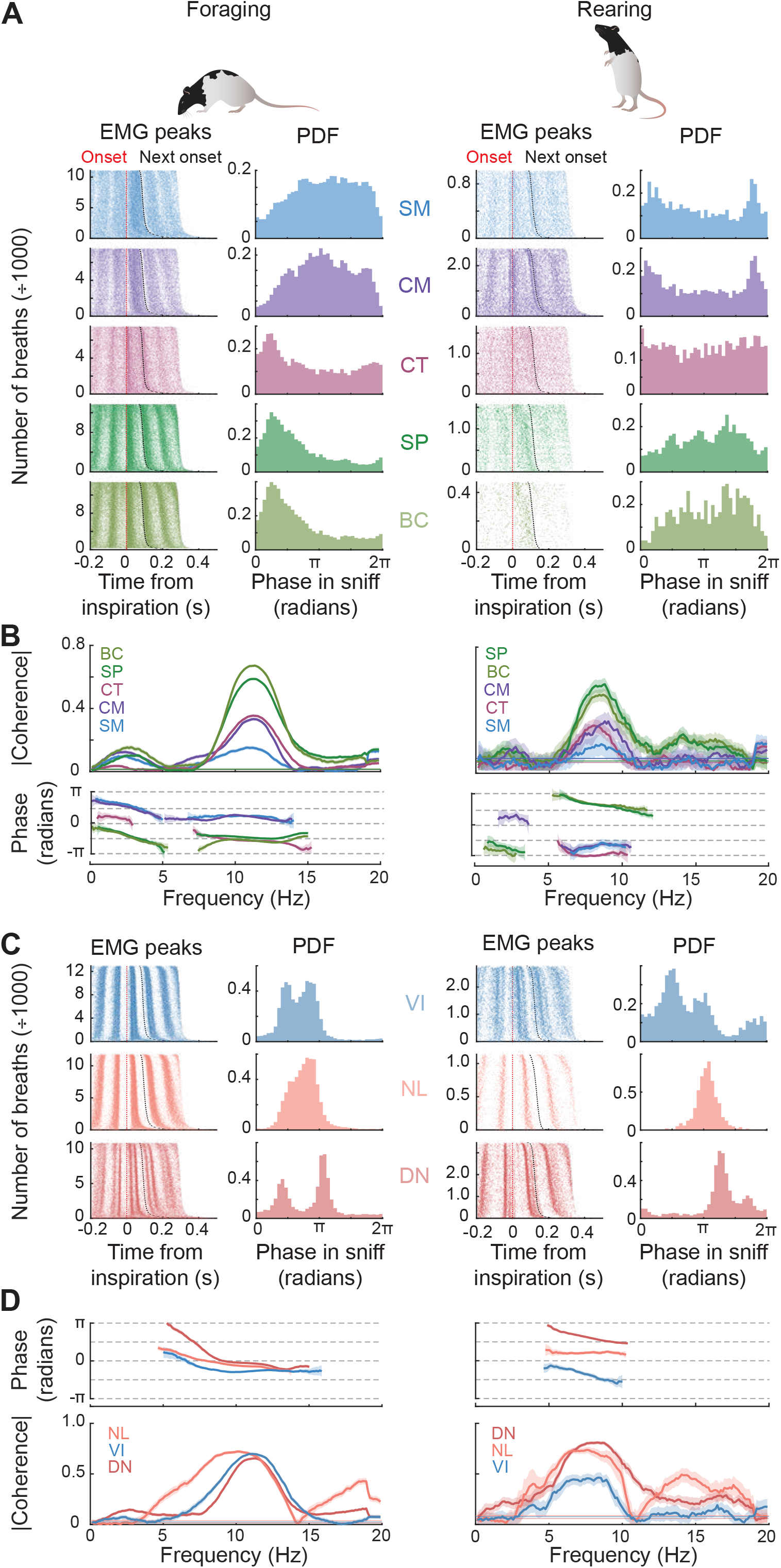
Quantification of the phase relation of neck and orofacial muscles to phase in the sniff cycle. **(A)** Raster plots of the neck EMG peaks with respect to the inspiration onsets (first and third columns), and the probability density functions (PDFs) of the neck EMG peaks in the sniffing cycles (second and fourth columns) during foraging (left 2 columns) and rearing (right 2 columns). Data shown from individual rats. **(B)** Coherence of neck EMGs with breathing in the foraging (left) and rearing (right) states. Coherence calculated from segments of length 4 s, half-bandwidth of 1 Hz (7 tapers). Shaded areas are the standard errors found with a jackknife procedure. Horizontal lines indicate the 0.95 confidence levels. Foraging data: SM (1,803 segments from 7 rats), CM (2,039 segments from 8 rats), CT (3,013 segments from 11 rats), SP (2,557 segments from 10 rats), BC (1,435 segments from 7 rats). Rearing data: SM (67 segments from 7 rats), CM (74 segments from 8 rats), CT (158 segments from 11 rats), SP (110 segments from 10 rats), BC (115 segments from 7 rats). **(C)** Raster plots of EMG peaks for vibrissa and nose movements with respect to the inspiration onsets, and the probability density function of EMG peaks in the sniffing (4 – 14 Hz) cycles. Data shown from individual rats. **(D)** Coherence of vibrissa and nose EMGs with breathing; parameters as in panel B. Data pooled from multiple rats (VI: 5, NL: 3, DN: 7). Shaded areas indicate the standard errors (jackknife). Horizontal lines show the 95% confidence levels. Number of segmented windows (foraging/rearing): VI (648/72), NL (288/55), and DN (1,668/122).

An alternate means to analyze the relation of muscle activity and breathing is in terms of spectral analysis. All five pairs of neck muscles were significantly modulated by breathing during both foraging, at 8 to 14 Hz, and rearing, at 6 to 11 Hz (**Figure 5B**); the range of frequencies for significance is consistent with the observed sniffing rate (**Figures 1H** and **5B**). Four of the five neck muscles, i.e., the sternomastoid, cleidomastoid, splenius, and biventer cervicis, shifted their phases with respect to breathing by ∼ π radians as the animal switched between foraging and rearing (**Figure 5B**). The exception is the clavotrapezius muscle, which is in phase with the splenius and biventer cervicis muscles during foraging, but only shifted its phase relationship with breathing by ∼ π/2 radians when the animal switched to rearing. Thus the phase relationship of the clavotrapezius muscle with respect to sniffing lies between the inspiratory, i.e., sternomastoid and cleidomastoid, and expiratory, i.e., splenius and biventer cervicis, subgroups of muscles during rearing (**Figure 5B**).

We now turn to a similar analysis for motor control of the vibrissae, mystacial pad, and nose. During foraging, the intrinsic vibrissa muscles (4 bilaterally and 1 unilaterally implanted rat) are activated at the start of inspiration, with the retractor nasolabialis (3 bilateral) activated after a delay (**Figure 5C**). Interestingly, the EMG envelope of the deflector nasi muscle (3 bilateral and 4 unilateral) consists of two peaks, one in the middle and one at the end of inspiration; this “double pump” was missed in the relatively sparse data set of prior work.^14^ The detailed pattern of activation shifted during rearing. A small change occurred for the intrinsic muscles, which are activated closer to the onset of inspiration, and for the retractor nasolabialis, which is activated later in the sniff-cycle. A large change occurred for the deflector nasi muscle, which is now activated during expiration (**Figure 5C**). Surprisingly, the activation patterns of these underlying muscles resemble the findings previously reported for head-fixed animals.^19,35,36,41^

The peak activity of each muscle group as a function of phase in the sniff-cycle was found from the spectral coherence (**Figure 5D**), similar to the case for neck muscles. The phase shifts for the intrinsic and nasolabialis muscles are small, about π/4 radians, between foraging and rearing. In contrast, the deflector nasi shifts its phase relationship with breathing by ∼ 0.65π radians when the animal switches to rearing (**Figure 5D**).

To further explore the dynamics of nose movements (3 rats), we formed a raster plot of the right deflector nasi muscle peaks with respect to the left peaks (**Figure S2A**), as well as their coherence (**Figure S2B**). Bilateral coactivation of deflector nasi muscle predominates under all conditions, yet unilateral activation is more prevalent during foraging than during rearing (**Figure S2A**,**B**). Thus, the rat performed more extensive lateral nose movements during foraging. Further to this point, nose twitching was phase-locked to head movement during both foraging and rearing (**Figure S2C**). Yet activation of the defector nasi muscle lags activation of the clavotrapezius muscle during foraging, but precedes activation of the clavotrapezius muscle during rearing. The most salient observation is that the vibrissa intrinsic and the deflector nasi muscles on the side of the head ipsilateral to a head-turn have greater magnitudes than those on the contralateral side (**Figure S2D**). This confirms that head-turning is accompanied by asymmetric whisking, with the ipsilateral vibrissae more protracted.^42,43^ From the perspective of behavior, these observations imply that the nose is drawn to the maximum lateral extent of a turn during foraging, consistent with efficient localization of an odor source.^35^

### Inspiration occurs at the transition from outward to inward horizontal turns

We have shown that egocentric, head-torso movements are phase-locked with breathing (**Figure 2A-C**). Yet horizontal head movements must also respect the midline, i.e., the symmetry axis of the body. To study the potential relationship between the timing of head-torso yaw movements and breathing relative to the midline, we separately analyzed the cases of egocentric counterclockwise and clockwise head-rotations during foraging. We identified the peaks in the head-torso yaw velocity for both directions of rotations (**Figure 6A**) and plotted the locations of the peaks with respect to the onset of inspiration as a raster (**Figure 6B**). For both counterclockwise and clockwise head-turns, the location of the peaks forms two clusters within the spectral band of sniffing. One cluster occurs during the inspiratory phase of a head-turn and the other during the expiratory phase (**Figure 6B**). We denote the four clusters as inspiratory counterclockwise (INS-CCW) and inspiratory clockwise (INS-CW) head-turns and as expiratory counterclockwise (EXP-CCW) and expiratory clockwise (EXP-CW) head-turns (**Figure 6C**).

**Figure 6.**
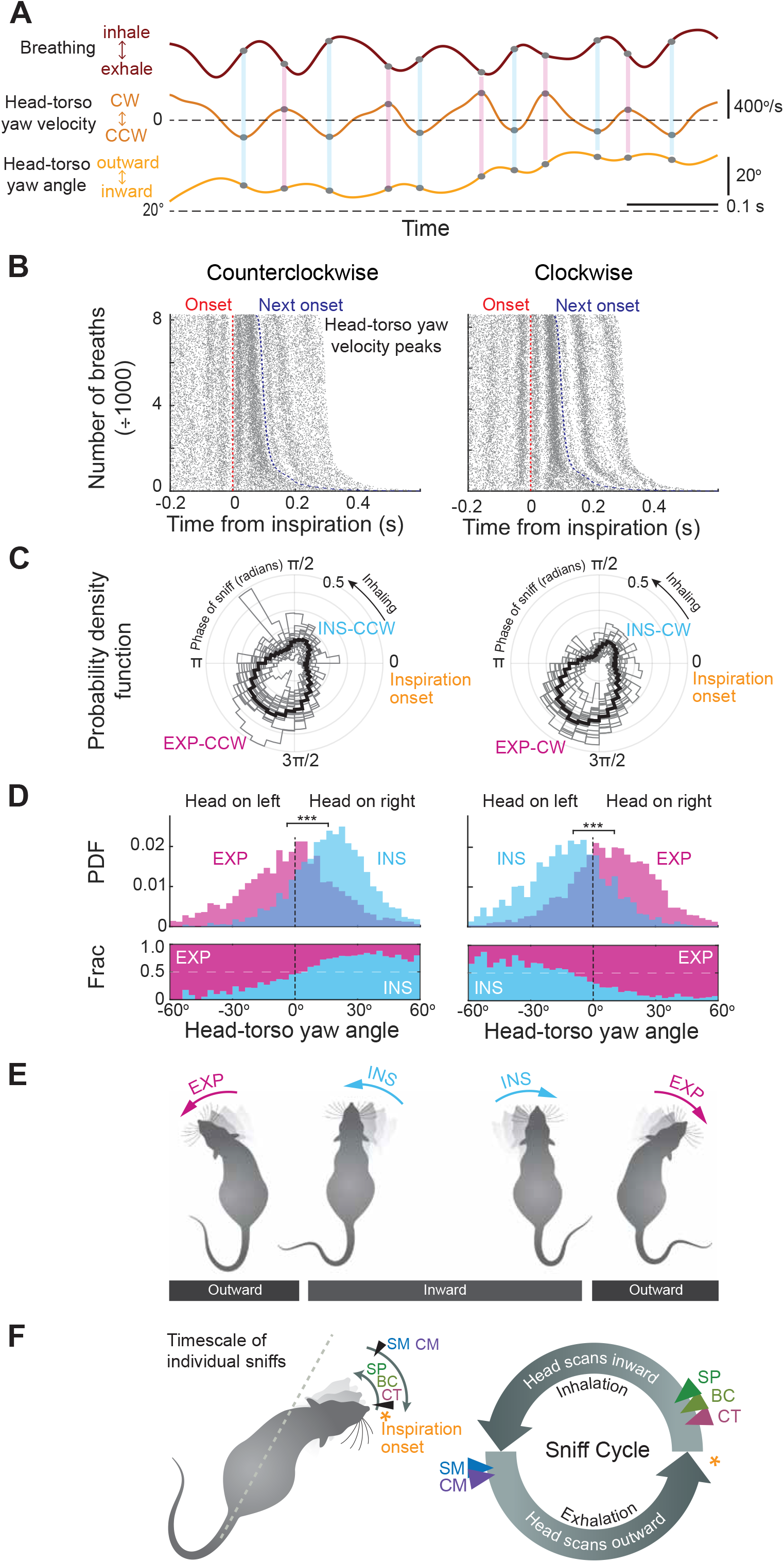
Inbound versus outbound horizontal head movements are preferentially biased to different phases in the breathing cycle. **(A)** Recording of breathing, head-torso yaw velocity, and head-torso yaw. Positive (clockwise) and negative (counterclockwise) peaks in the velocity data are marked. **(B)** Raster plots of counterclockwise (CCW, left) and clockwise (CW, right) head-torso yaw velocity peaks with respect to the inspiration onsets. Rows are sorted by the periods of the breathing cycles. Red dashed line indicates the onsets of the breathing, and black dashed line indicates the subsequent inspiration onsets. Data from one rat. **(C)** Probability density function of the sniffing phase where the head-torso yaw velocity peaks (left: CCW, right: CW) occur, plotted in polar coordinates (17 rats). Black curve shows the results combined from all 17 rats. Windows of size π/2 radians centered at 0.96 radians and 4.28 radians define the regions of inspiratory (INS) and expiratory (EXP) head movements. **(D)** Probability density function (upper) of the instantaneous head-torso yaw angle associated with inspiratory and expiratory head-wiggles (left = CCW; right = CW) and their fractional proportions (lower). Data from one rat. Head-torso yaw angle is defined to be positive (or negative) when the head is on the right (or left) of the torso midline. **(E)** Depiction of the postural dependency of the bimodal oscillations of the head. For both CW and CCW rotations, inward turns are more likely to occur with inspiratory breathing, while outward turns are more likely to occur with expiratory breathing. **(F)** Depiction of head scanning with sequential activation of the neck muscles in the sniff cycle.

We found that the division of inspiration and expiration events depends on the location of the head relative to the midline of the animal (p < 0.001 in 16 of 17 rats, Kolmogorov-Smirnov test) (**Figure 6D**). Inward turns, i.e., toward the midline, are more likely to occur during inspiration for both counterclockwise and clockwise head rotations (**Figure 6E**). As such, the initiation of outward turns, i.e., away from the midline, are more likely to occur during expiration (**Figure 6E**). Thus the next inspiratory event during an outward turn occurs with the head at maximum rotation. This ensures a fresh sampling of the environment.

We summarized the sequence of neck muscle recruitments during the sniffing cycles by combining the data for movement (**Figure 6C**,**D**) with the EMG recordings from neck muscles (**Figure 5A**,**B**). Upon the onset of inspiration, the clavotrapezius, biventer cervicis, and splenius muscles are recruited to turn the head inward (**Figure 6F**). At the transition from inhalation to exhalation, the sternomastoid and cleidomastoid muscles are recruited and turn the head outward (**Figure 6F**). The inward versus outward dependence of horizontal head movement with sniffing is less evident when the rat is in the rearing state (p = 0.0013 for CCW and p = 0.16 for CW head movements; Kolmogorov-Smirnov test) (**Figure S3A**,**B**).

Other features may contribute to the division of inspiratory versus expiratory head-turns. We find a bimodal distribution for the radial location in the arena (p < 0.01 in 5 of 17 rats, Kolmogorov-Smirnov test) (**Figure S3C**,**D**), for the locomotion speed (p < 0.001 in 11 of 17 rats, Kolmogorov-Smirnov test) (**Figure S3E**), and for the instantaneous speed of the head-turn (p < 0.001 in 16 of 17 rats, Kolmogorov-Smirnov test) (**Figure S3F**). In general, the maximal head-speed during expiratory turns is greater than the maximal speed in inspiratory turns (**Figure S3F**).

The activation of all five neck muscles relative to breathing has a peak at zero lag time (**Figures S1D** and **S3G**). This indicates that bilateral muscles are coactivated, as occurs at most joints^44^, and the net torque determines the resultant direction of the head-rotation. Yet the cleidomastoid muscle, unlike the other four neck muscles, showed peaks at ± 0.25 s in the cross-correlation between its left and right instantiations (**Figure S3G**); this is in addition to its participation in synergistic control of head-turning (**Figures 3** and **5**). Thus the cleidomastoid muscle is also responsible for controlling the slow component of the horizontal head orientation (**Figure 2B**), during which the EMG envelope of the ipsilateral cleidomastoid muscle exhibits a sequence of peaks that correspond to small, successive head rotations across several sniffs (**Figure S3H**).

## DISCUSSION

Our first claim involves the modulation of breathing, the fundamental rhythm of life. When rodents switch from foraging, with their vibrissae sweeping across the ground and their nose scanning the air just above, to the state of rearing, when they rise on their hind legs, their breathing rate decrements by 3 Hz (**Figure 1H**). Changes in posture could lead to a change in breathing rate based solely on pulmonary mechanics. Yet, breathing may serve as a proxy for behavioral state that is readily decoded into movement and action.^26^ One possibility is via activation of locus coeruleus by breathing.^45^

Our second claim is that the activation of neck and orofacial muscles relative to breathing depends on behavioral state (**Figures 5** and **7A**). During foraging, the clavotrapezius, splenius, and biventer cervicis muscles are activated during inspiration, while the sternomastoid and cleidomastoid are recruited during expiration. These muscle synergies lead to an outward head rotation during expiration and an inward head rotation at the onset of inspiration. The switch from foraging to rearing is marked by an inversion of this organization, in which the phase-shift with respect to breathing is nearly π radians while the phase difference between muscle groups is maintained (**Figures 5B** and **7A**). The one exception is the clavotrapezius (**Figure 5B**). From the perspective of modulating sensation by motor output, it may be most reliable to issue commands to modulate separate pools of motoneurons that control head movement by binding them to a common rhythm. All told, our results revealed a previously unrealized computational complexity that emerges from hindbrain circuits.

Our third claim is that the musculature involved in head-turning and nose turning coordinate with breathing in a manner that respects the midline as an axis of symmetry (**Figure 6E**,**F**). The rat tends to turn outward for both counterclockwise and clockwise rotations during expiration, and turn inward toward the midline after the onset of inspiration. The observations that head-yaw rotation during expiration has a greater average angular speed than rotation upon inspiration, and that the outward head rotation in egocentric coordinates occurs during expiration, are consistent with the notion that the expiratory phase of breathing is used to relocate the motor plant for a subsequent “snapshot” sample of the environment.^14,17,24^ Recalling that the rat also uses the expiratory phase to relocate the head vertically^14^, the motor strategy of head movement is similar for both the vertical (pitch) and horizontal (yaw) axes.

Our work highlights the necessity for discovery based on the use of freely moving animals. In the current paradigm, the sniffing rate lies between 8 and 14 Hz (**Figure 1H**), as compared with a substantially lower rate of 4 to 8 Hz in studies with head-fixed rats.^14,17^ The use of a large arena allowed the animal to readily change pose, with accompanying changes in breathing rate. A similar change in the rate of sniffing with a change in task occurs with rodents that sample an odor and then change state to respond with a head-poke and potentially receive a reward.^30^ More complicated behavioral paradigms may lead to rodents taking on additional poses and reveal further complexity in the relation of motor actions to breathing.

### Phase shift among behavioral states

Two changes occur as the rat alters its behavioral state from foraging to rearing. First, the central breathing frequency shifts from 11 to 8 Hz (**Figure 1H**). Second, the sternomastoid and cleidomastoid muscles shift their phase relationship with sniffing from expiratory to inspiratory activity, while the splenius and biventer cervicis muscles shift their phase relationship with sniffing from inspiratory to expiratory activity (**Figures 3A** and **5B**). This combination is reminiscent of the coordinated shift in locomotion frequency and gait in tetrapod.^6,8,9,11^ It may be modeled by coupled phase oscillators^46^ in which the spinal segments are taken as oscillators^12,47^ and the interactions include time lags^46,48^.

Our model for coupling of neck movement to breathing has three oscillators (**Figure 7B**). The oscillator for inhallation, the preBötzinger complex (pBötC)^49,50^, has a variable output frequency that is taken as either *f*_pBötC_ = *f*_forage_ = 11 Hz for the foraging state or *f*_pBötC_ = *f*_rear_ = 8 Hz for the rearing state (**Figure 7B**). Two oscillators, connected with reciprocal, instantaneous inhibitory connections, represent the neck oscillators. Each neck oscillator drives one of the two antiphase subpopulations of neck motoneurons. The neck oscillators have an intrinsic frequency that is the mean value of *f*_pBötC_, or *f*_neck_ = (*f*_forage_ + *f*_rear_)/2 = 9.5 Hz. A unilateral connection from the pBötC to one of the two neck oscillators has strength Γ_B_ and a time lag of τ_neck_. We show (Star Methods) that there are pairs of values for Γ_B_ and τ_neck_ for which the neck oscillator will shift in phase by π radians relative to the breathing oscillator when the frequency of sniffing changes between foraging and rearing. A delay of τ_neck_ = 20 ms is mathematically feasible and is biologically plausible in terms of slow adaptation currents (**Figure S4A**). Further, the - 0.75π radians phase offset between the two subgroups of neck oscillators is incorporated through the choice of the inhibitory connection strength.

**Figure 7.**
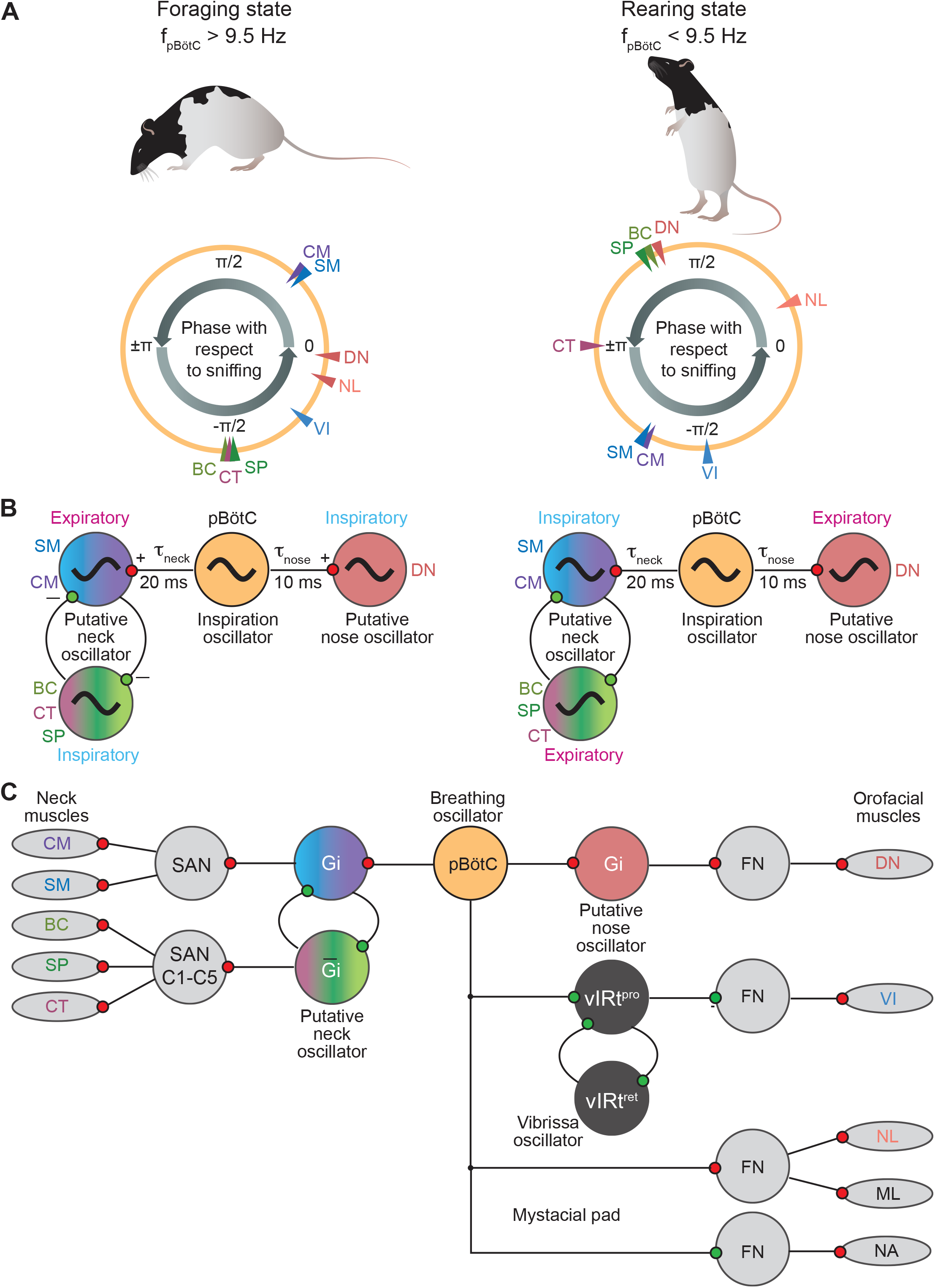
Summary of phase shifts and models. **(A)** Depiction of the phase relationship of neck and orofacial muscles with respect to breathing. Phase is defined from the coherence at the sniffing frequency (**Figure 5B,D**). Note the large changes for neck muscles, more modest change for the nose muscle (DN), and small changes for the whisking and mystacial muscles during foraging versus rearing. **(B)** Illustration of the interaction between breathing and head and nose movements using unidirectional coupling oscillators models. Red indicates excitatory synapses and green indicates inhibitory synapses in our model (Star Methods). **(C)** Cartoon of the known circuit for the control of rhythmic whisking ^22^ and potential circuit for the medullary control of head-turning and nose movements ^67,68^. New abbreviations: SAN = spinal accessory nucleus; C1-C5 = spinal motonuclei at cervical levels 1 to 5; Gi = Gigantocellular nucleus in the medulla; FN = facial motonucleus; vIRt^pro^ and vIRt^ret^ = protraction and retraction subregions of the vibrissa premotor oscillator; ML and NA = maxillolabialis and nasalis muscles of the mystacial pad.

The deflector nasi also shows a shift in coordination that depends on foraging versus rearing (**Figure 5D**). Premotor neurons of deflector nasi have been identified in the reticular formation, and some receive the projections from the pBötC.^35^ Thus a similar analysis, except for only two oscillators, show that a frequency-dependent phase-locking holds for sniffing and nose wiggling (**Figure 7B**). Here, the pair with a shift in phase of 0.65π radians can be achieved with τ_nose_ = 10 ms (**Figure S4B**).

All told, our analysis shows how, in principle, driving the sniffing rate at different frequencies will allow the rat to entrain the neck muscles at different phases in the sniff cycle to fulfill a specific behavioral need. There is no need for plastic changes in connectivity in the brainstem circuitry. The phase shifts can presumably be fine-tuned by neurobiological realism.^51^

### Potential underlying neuronal circuit

The hierarchical arrangement that governs the impact of breathing on whisking was delimited over the past decade.^17,19,20,22^ Two populations of inhibitory neurons in the vibrissa intermediate reticular formation form a network oscillator that can free run or be paced by projections from the pBötC.^49^ Neighboring regions in the intermediate reticular formation likely contain putative oscillator circuits for the orofacial circuits that drive the muscles involved in licking^52-56^, chewing^55,57-59^, and nose twitching.^35^

The distributions of the motoneuron pools of the five neck muscles we examined (**Figure 3A**) have been well studied^60,61^, yet their premotor nuclei have not been fully investigated. It has been reported that there are no apparent direct projections from the pBötC to the spinal cord.^62^ Therefore, the pathway of the respiratory drive for the neck muscles should involve premotor circuitry on the medulla, although relays of the respiratory oscillators^62,63^ or the high cervical respiratory group in the cervical spinal cord^64,65^ cannot be discounted.

Retrograde tracing with Cholera toxin subunit B from sternomastoid revealed labeled neurons in the reticular formation.^66^ Subsequent viral tracing studies localized the gigantocellular region of the reticular formation as a prospective location of neck premotor neurons, most likely V2a neurons.^67,68^ These cells receive descending input from the contralateral superior colliculus, a high-level region that drives orienting and thus head position in rats.^69,70^ In addition to the reticular formation, coordination between neck motoneurons may involve spinal interneurons in the propriospinal networks^71^, especially as microstimulation from vertebral levels C2 to T1 elicits sternomastoid activity.^72^ In addition to connections from the pBötC to the gigantocellular premotor neurons for movement of the neck, there is some evidence for a similar pathway for rhythmic movement of the nose.^67^ These potential connections, along with known, related connections for rhythmic movement of the vibrissa and the mystacial pad^17,19,20^, are summarized in **Figure 7C**.

### Potential feedback control

Past studies identified the participation of sensory feedback in the control of whisking. The anatomical pathway involves vibrissa input to the spinal trigeminal subnuclei rostral interpolaris and oralis^73,74^ that, in turn, project to vibrissa and mystacial motoneurons in the facial motor nucleus to complete a vibrissa-trigemino-facial reflex.^73-75^ It is noteworthy that the prolonged, flat-top response as the rat’s vibrissae contact the floor of the arena during foraging (* in **Figure 4D**), reminiscent of touch-induced “pump”^76,77^ may be explained by a transient inhibitory reflex.^73^ Lastly, neurons in spinal trigeminal oralis send projections to the cervical spinal cord.^78^ This might serve as the substrate for coordinated movements of the vibrissae and the head to ensure proper timing of contact of the vibrissae to the ground in the sniff cycle.^43,79-81^

## Supporting information

Supplemental Figres and Table

## ACKNOWLEDGEMENTS

We thank Rodolfo Figueroa for assistance with the instrumentation, Zaneta Ku, Peyton O’Callaghan, Emma Osgood, Sabrina Tuell and Nina Zhang for animal training, Julia Kuhl for drawing all illustrations, Jill Leutgeb, Fan Wang, Jing Wang and Arash Fassihi Zakeri for conversations, and Beth Friedman and Philipp Mächler for comments on a draft manuscript. This work was supported by the Taiwan Ministry of Education Scholarship and the United States National Institutes of Health (R01 NS058668; U01 NS090595; U19 NS107466).

## CONTRIBUTIONS

DK and S-ML conceived the project and codeveloped the experimental approach, the instrumentation, the data analysis procedures, and the model. S-ML performed all experiments and discovered the previously undescribed phase shifts with behavioral state. DK and S-ML co-wrote the manuscript. DK attended to the myriad of university compliance rules and workshops that govern the ethical use of animals, environmental health and safety as well as research integrity and ethics and the maintenance of a harassment-free work environment for all laboratory personnel.

## STAR Experimental Procedures

### KEY RESOURCES TABLE

| REAGENT or RESOURCE | SOURCE | IDENTIFIER |
| --- | --- | --- |
| Chemicals, peptides, and recombinant proteins |  |  |
| Isoflurane | Henry Schein | Cat no. 1182097 |
| Ketamine, Injection | Zoetis | Cat no. 40027676 |
| Xylazine, Injection | AnaSed | Cat no. (01)00359399110204 |
| Buprenorphine Hydrochloride, Injection | Par Pharmaceutical | Cat no. 110483373347 |
| Experimental models: Organisms/strains |  |  |
| Rat: Long-Evans | Charles River Laboratories |  |
| Software and algorithms |  |  |
| MATLAB | MathWorks |  |
| Chronux | Cold Spring Harbor Laboratory |  |
| DeepLabCut | https://github.com/DeepLabCut/DeepLabCut |  |
| LabChart | AD Instruments |  |
| Pylon | Basler |  |
| Arduino IDE | Arduino |  |
| Tennsyduino | https://www.pjrc.com/ |  |
| BNO Library | Adafruit |  |
| Other |  |  |
| Absolute Orientation Sensor | Adafruit | Cat no. BNO055 |
| Microcontroller | https://www.pjrc.com/ | Cat no. TEENSY3.2 |
| CCD camera | Basler | Cat no. acA2040-90umNIR |
| PowerLab | AD Instruments | Cat no. 16/35 |
| Differential amplifier (thermocouple) | World Precision Instruments | Cat no. DAM80 |
| EMG Preamplifier | Kleinfeld Laboratory | https://neurophysics.ucsd.edu/lab/16_channel_ephys_second_stage.pdf |
| Current stimulator | A-M Systems | Cat no. 2100 |
| Thermocouple | Omega | Cat no. 5TC-TT-K-36-36 |
| Tungsten wire for electromyographs | A-M Systems | Cat no. 795500 |
| 10-pin connector | Samtec | Cat no. TMS-110-01-G-S-RA (M)<br>Cat no. SMS-110-01-G-S (F) and SMS-110-01-L-S (F) |

## RESOURCE AVAILABILITY

### Lead contact

Further information and requests for resources and reagents should be directed to and will be fulfilled by Prof. David Kleinfeld.

### Materials availability

N/A

### Data and code availability

The datasets supporting the current study, and an associated “read me” file, will be available at https://datadryad.org/ upon publication. The code for the model will be available upon publication at https://github.com/.

## EXPERIMENTAL MODEL AND SUBJECT DETAILS

### Experimental subjects

We used 33 Long Evans adult female rats ranging from 240 to 430 grams for this study. Behavioral training and surgical procedures were in accordance with the animal use protocol approved by the Institutional Animal Care and Use Committee (IACUC) at the University of California San Diego.

### Behavioral training

Animals (Long-Evans rats) were trained to forage for food under food restrictions. Before food restriction, body weight was measured to establish a baseline. Each food restriction period was no longer than five consecutive days. After five consecutive days of food restriction, animals had at least two days of ad libitum access to food.

Animals were weighed daily during the period of food restriction. Restriction ended immediately if the weight dropped below 0.8-times baseline. During the food restriction period, supplemental food was provided to ensure that the rats were not deprived of food for more than 24 hours.

During the training and recording sessions, the rat was placed inside the foraging arena. Food pellets of ∼ 0.1 g were dropped into the arena one at a time at random locations after an auditory cue. The experiment ended when the rat no longer foraged for food.

### Surgical procedures

All procedures were performed under anesthesia through the injection of ketamine (50 mg/kg-rat) and xylazine (5 mg/kg-rat). The level of anesthesia was monitored regularly by pinching the foot pad. Supplemental doses of ketamine (15 mg/kg-rat) and xylazine (1.5 mg/kg-rat) were provided when needed. An injection of buprenorphine (0.03-0.05 mg/kg-rat) was given before and after the surgery. At surgery, an incision was made along the midline above the skull to the nose. After cleaning the skull surface, 6 to 8 no. 00-90 screws (McMaster-Carr) were implanted. A hole (Bur Carbide FG ½, Henry Schein) was drilled on top of the nasal cavity, and a sterile thermocouple (5TC-TT-K-36-36, Omega) was inserted into the nasal cavity. The thermocouple wires were attached to the skull with Loctite 401 and Jet Acrylic (Lang Dental), and the end was soldered to a pair of 2.5 mm male pin connectors.

The electromyogram (EMG) electrodes were made with tungsten wires (#795500, A-M Systems). A pair of tungsten wires, with insulation removed at ∼1 mm at the tips, were aligned with a 1 to 3 mm distance between the bare tips, depending on the size of the target muscle. The tips of the electrodes were bent to create a hook. For EMG recordings in intrinsic vibrissae muscles, bipolar needle electrodes^82^ were used, where the electrodes are passed through a 25-26G hypodermic needle. For EMG recordings in all other muscles, bipolar suture electrodes^82^ were used, where the hook of the tungsten electrodes was tied to a silk suture. Electrodes were autoclaved before surgery. For implanting electrodes to the vibrissa intrinsic muscles, the hypodermic needle was passed subcutaneously from above the nasal bone to reach the vibrissae follicles. The needle was then gently retracted, leaving the electrodes in place. For implanting electrodes into all other muscles, surgical procedures were performed to expose the target muscle. Target muscles were identified using literature as references^34,36,40,60,61^ and dissection studies (15 rats). The deflector nasi and nasolabialis muscles were accessed from above the nasal bone. Access from the dorsal neck was made to expose muscles splenius, biventer cervicis, and clavotrapezius. Access from the ventral neck was made to expose muscles sternomastoid and cleidomastoid. After exposing the target muscle, a silk suture was passed through the muscle belly gently to drag the electrodes into the muscle body. After exiting, the silk was tied back to the facia near the entry point to secure the location of the electrodes. We used electrical stimulation (Model 2100, A-M Systems) to confirm the location of the implanted electrodes by sending a pulse train of 0.2 ms duration, 4.8 ms burst width, and 1.2 ms inter-pulse period with 100 - 500 μA of current ^36^. A reference electrode was made with a tungsten wire (#795500, A-M Systems), with ∼ 5 mm striped from the tip. The reference electrode was placed under the skin near the incision. The ground wire was made of a silver wire (#786000, A-M Systems) soldered to one of the head screws. Finally, the ends of the tungsten and silver electrodes were soldered to 10-pin male connectors (Samtec). All surgical incisions were closed with sutures.

In 17 animals, an orientation sensor was implanted into the torso subcutaneously between T1-T6 vertebrae by incision on the back. The torso sensor (BNO055, Adafruit) was covered with epoxy for insulation and sterilized. The surgical incision was closed with sutures. In all animals, a 10-pin female connector (Samtec) was fixed to the skull with Jet Acrylic. The head orientation sensor (BNO055, Adafruit) was attached to the connector before the start of each recording session. After surgery, animals were allowed to rest for at least two full days. Post-operative animals were checked regularly to monitor their conditions. Details of the procedures of each animal are listed in **Table S1**.

## METHOD DETAILS

### Video annotation and location tracking

As the rat was searching inside the arena, the entire process was recorded through a Basler camera (acA2040-90umNIR) above the arena at 20 fps. We used DeepLabCut^83,84^, a 2-D convolutional neural network-based algorithm, to track the animal’s location. From each recording video, about 20 frames picked by DeepLabCut were manually labeled to mark the location of the rat’s lower (sacral) torso. The labeled data were split into a 19:1 ratio for training and validation. We used a batch size of 1 and a learning rate of 0.005 with the SGD optimizer and trained a Resnet-50 model^85,86^ for two iterations (20,000 epochs each). Other parameters were set to default. The tracking results were saved as a CSV file. To locate the center of the arena, we fitted the image of the floor boundary with an ellipse^87^.

We inspected all video files to manually in order to label the frames where the rat performed miscellaneous behaviors, including scratching, dog-shaking, grooming, urinating, defecating, biting, and freezing. Data taken that these miscellaneous behaviors were not used in the analyses.

### Data recording and pre-processing

The breathing signal was recorded from the rat by connecting the thermocouple to an amplifier (DAM80, World Precision Instruments). We used a 0.1 Hz high-pass filter, 100 Hz low-pass filter, and 10,000-times gain. The EMG signals were connected to a pre-amplification stage of local design to obtain a gain of ×400 and were high pass filtered at 0.1 Hz (https://neurophysics.ucsd.edu/lab/16_channel_ephys_second_stage.pdf). Breathing and EMG signals were sampled either at 20 kHz (27 rats) or 40 kHz (6 rats) with the data acquisition system (PowerLab, ADInstruments).

Head and torso orientation sensors (BNO055, Adafruit) were connected to a development board (Teensy 3.2, PJRC) via the I2C port. The orientation and movement signals from the head and torso sensors were read by Arduino code adopted from the Adafruit BNO055 library (https://github.com/adafruit/Adafruit_BNO055). Signals were sampled at 100 Hz, and the timestamps of each sample were sent to the breathing and EMG acquisition system with a pulse signal. Sensor data were displayed on the Arduino Serial Monitor and were saved to the hard drive at intervals no longer 7 minutes, and typically 6-½ minutes, at which time the head and torso sensors offsets drifted by less than one resolution unit or 0.01°.

Pre-processing of data was done in MATLAB (MathWorks). First, we took the difference in the EMG signals between the pair of electrodesb to obtain the differential EMG. The differential EMG was low passed at 300 Hz by a third order Butterworth low-pass filter and high passed at 9,999 Hz by a third order Butterworth high-pass filter. We then obtained the demodulated EMG envelope by taking the absolute value of the signal, low passed at 50 Hz with a third-order Butterworth filter, and down-sampled to 2 kHz.

Digitized breathing data were low passed at 20 Hz with a fifth-order Butterworth filter, high passed at 1 Hz with a third-order Butterworth filter, and down-sampled to 2 kHz. The torso location-tracking data were low passed at 4 Hz with a third-order Butterworth filter and were interpolated to 2 kHz with a cubic spline. The head and torso orientation data were interpolated to 20 kHz with a cubic spline. The head orientation data were low passed at 25 Hz with a third-order Butterworth filter. The torso orientation data were low passed at 4 Hz with a third-order Butterworth filter. All digital filters were run in both forward and reverse directions to obtain zero phase distortion. Finally, we took the difference between the head yaw and torso yaw to obtain the relative head-torso yaw, and we set the mean of the head-torso yaw angle of the entire recording to zero.

## MODEL

### Two-cell circuit

We analyze a dynamical system comprised of two phase-oscillators to model rhythmic nose twitching. We assume the connection between the two oscillators is unidirectional ^88^, where only the nose oscillator is receiving signals from the breathing oscillator but not vice versa (**Figure 7B**). We consider a time lag constant *τ* and write

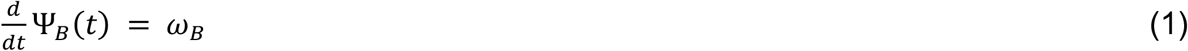

and

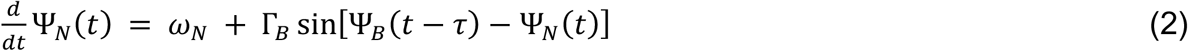

where Ψ_B_(t) is the phase of the breathing oscillator, in units of radians, and Ψ_N_(t) is the phase of the presumed nose oscillator. The coupling strength is denoted by Γ_B_, the intrinsic frequency of the nose oscillator is denoted by ω_N_, and the intrinsic frequency of the breathing oscillator is denoted by ω_B_; all coupling strengths and frequencies are in units of rad/s. When two oscillators are phase-locked, we have

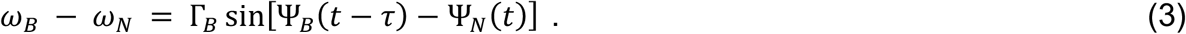

We make the ansatz that the solution takes the form of two oscillators that are locked at the breathing frequency, with a phase difference *α*. Thus

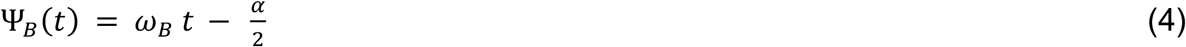

and

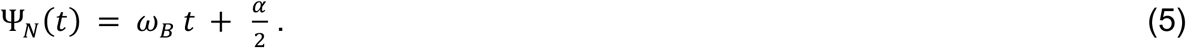

Plugging Eqs (4) and (5) into Eq (3) yields:

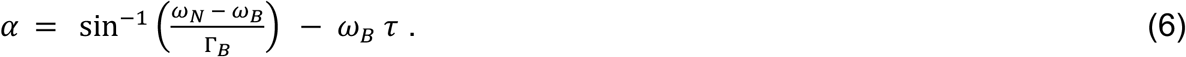

The phase α corresponds to the phase between the breathing and nose oscillators and depends on two unknowns, Γ_B_ and τ. The solution exists, i.e., the breathing oscillator can entrain the nose oscillator, when

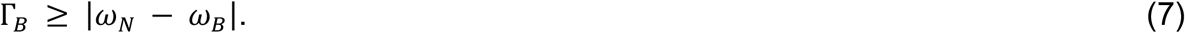

#### Stability analysis

The stability of the solutions in the model can be determined by adding a small perturbation to the steady-state solution of the nose oscillator, i.e.,

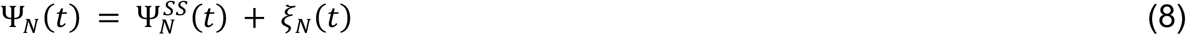

Substituting the right-hand-side of Eq (8), into Eqs (2), (4) and (5) yields

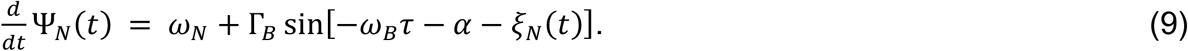

We expand the above equation for small magnitudes of *ξ*_*N*_(*t*) to:

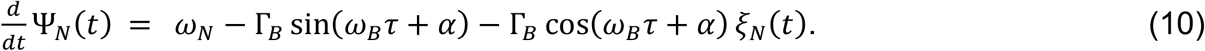

Since

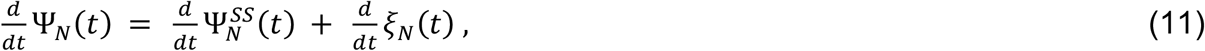

the steady state solution will factor out of Eq (10) and we obtain an equation for the perturbation, i.e.,

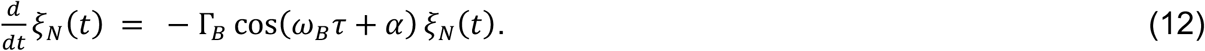

Substituting in Eq (6) yields

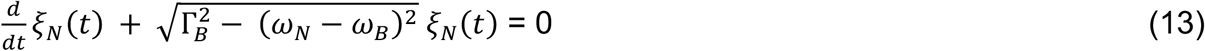

and leads to a stable system, i.e., *ξ*_*N*_(*t*) decays to zero, by the existence criteria of Eq (7) for both foraging and rearing (**Figure S4**). Note that the recovery rate increases as the coupling strength increases.

### Three-cell circuit

We analyze a dynamical system comprised of three phase oscillators to model rhythmic neck rotations. We assume a unilateral connection from the breathing oscillator to one of the two neck oscillators and reciprocal connections between the two neck oscillators (**Figure 7B**). We consider a time lag constant *τ* only for the unilateral connection and write

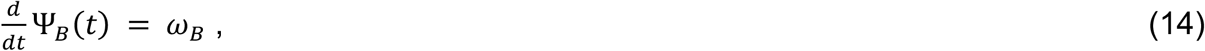

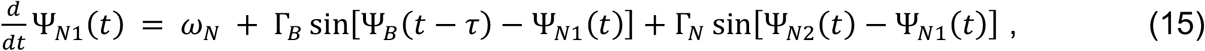

and

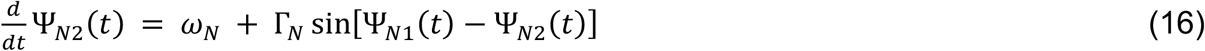

where the reciprocal coupling strength between neck muscles is denoted by Γ_N_, and the intrinsic frequency of the neck oscillator is denoted by ω_N_. When three oscillators are phase-locked, we have

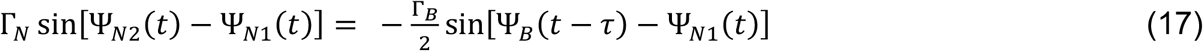

and

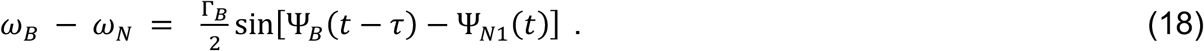

Similar to the case for two oscillators, we make the ansatz that the solution takes the form of three oscillators that are locked at the breathing frequency, with

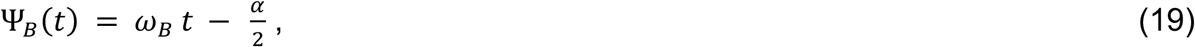

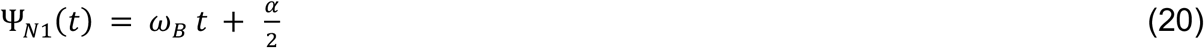

and

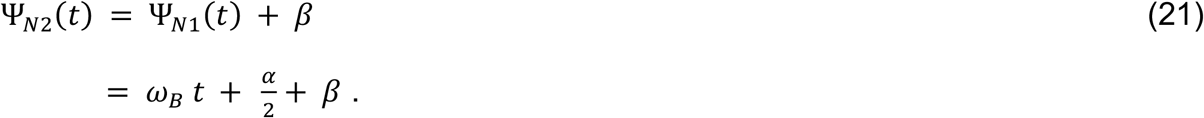

Plugging Eqs (19) to (21) into Eqs (17) and (18) yields:

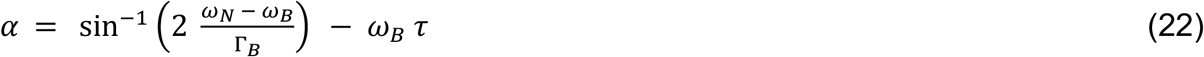

and

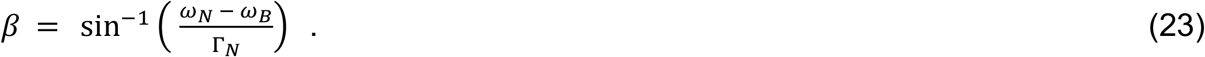

The phase α corresponds to the phase between the breathing and one population of neck oscillators and depends on two unknowns, Γ_B_ and τ. The phase β corresponds to the phase between the two populations of neck oscillators and depends on one unknown, Γ_N_.

The solution exists when two criteria are satisfied. First, when:

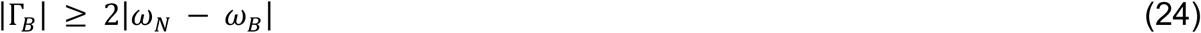

and second, when

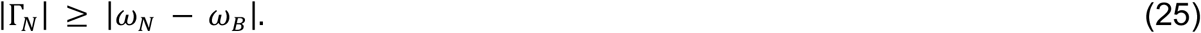

#### Stability analysis

The analysis of the three-cell circuit divided into that of two pair-wise circuits. The stability of the breathing oscillator and nose oscillator pair was discussed previously. For the pair of neck oscillators, stability can be also determined by adding a small perturbation to the steady-state solution of the neck oscillators, i.e.,

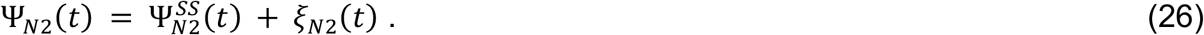

Substituting the right-hand-side of Eq (26), into Eq (16) yields

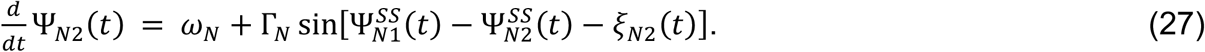

We expand the above equation for small magnitudes of *ξ*_*N*2_(*t*) and factor out the steady state response to obtain an equation for the perturbation, i.e.,

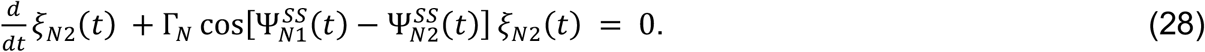

Or

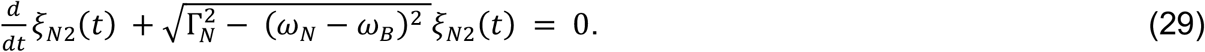

This leads to a stable system, i.e., *ξ*_*N*2_ (*t*) decays to zero, when Eq (25) is satisfied. Further, phase shifts between Ψ_N1_(t) and Ψ_N2_(t) within the neighborhood of β = π radians are achieved, by Eq (28), with Γ_N_ < 0. This implies mutual inhibition, for which π/2 < β < 3π/2, with β → π as Γ_N_ → -∞

### Application

The shift in the phase between foraging and rearing corresponds to the shift in α between these two states. We assume that ω_B_ = 22π rad/s (11 Hz) during foraging and ω_B_ = 16π rad/s (8 Hz) during rearing (**Figure 1F**). We further assume that all coupling strengths and the intrinsic frequency of all but the breathing oscillator are constants.

#### Breathing drives the neck oscillator

Or measurements of the shift in phase correspond to the difference α_*forage*_ - α_*rear*_. From Eq (22),

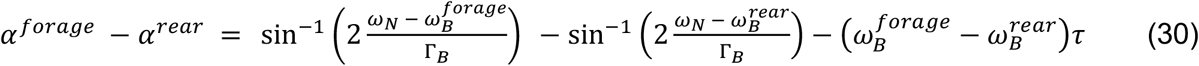

which is one equation in two unknowns, i.e., τ and Γ_B_. From the data, α^*forage*^ - α^*rear*^ = π (**Figure 5B**) and we assume ω_N_ = 19π rad/s (9.5 Hz), chosen as the midpoint frequency between rearing and foraging (**Figure 1H**). We solve Eq (30) numerically to find all pairs of values of τ and Γ_B_ (**Figure S4A**). From Eq (24), the magnitude of Γ_B_ is bounded by Γ_B_ β 6π rad/s. A plausible pair of solutions is τ = 20 ms and Γ_B_ = 19.2 rad/s.

The phase shift among the two neck oscillators is observed to be β = -0.75π **(Figure 5B**). We find Γ_N_ = -13.2 rad/s from Eq (23).

#### Breathing drives the nose oscillator

This analysis is similar to that for the neck except for a numerical difference in formula (Eq (6)). Thus

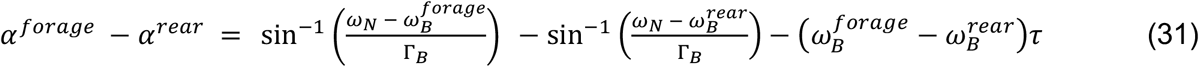

From the data, α^*forage*^ - α^*rear*^ = 0.65π (**Figure 5D**) and as above we assume ω_N_ = 19π rad/s (9.5 Hz). We solve Eq (31) numerically to find all pairs of values of τ and Γ_B_ (**Figure S4B**). From Eq (7), the magnitude of Γ_B_ is bounded by Γ_B_ ≥ 3π rad/s. A plausible pair of solutions is τ = 10 ms and Γ_B_ = 11.8 rad/s.

## DATA ANALYSIS

We used MATLAB (MathWorks) code for data analysis. Spectral analyses were performed with Chronix (http://chronux.org/).^89^ To define the inspiration onsets, we followed the procedures from a previous studies: the Hilbert transform was applied on the thermocouple signal to extract all the local peaks (maximal inhalation) and troughs (maximal exhalation)^41^, and the inspiration onsets were defined to be the 10 % rise times from each trough.^17^ The peaks in the movement signals, recorded from head or torso orientation sensor, were identified by setting the 75^th^ percentile value as the minimum height threshold and the 0.5 x std as the minimum prominence; percentiles and std were calculated from all recording sessions in the same rat. The peaks in the EMG envelopes were identified by setting the 90^th^ percentile value as the minimum height threshold, the 99.99^th^ percentile value as the maximum height threshold, and the 0.5 x std as the minimum prominence; percentiles and std were calculated from all recording sessions in the same rat.

