## Supplemental Figres and Table for "A change in behavioral state switches the pattern of motor output that underlies rhythmic head and orofacial movements"

**(G)** Cross-correlation of left and right neck EMG signals in the foraging state, averaged from segments of length 4 s. Data from individual rats (SM and CM: 34 segments, CT: 152 segments, SP and BC: 144 segments). Error bars are standard errors. Asterisks mark the activities related to head orientation.

**(H)** Illustration of the activation pattern of CM in head orientation.

**Figure S4: Computational models for phase shifts.**

**(A)** Unidirectional coupling model for head movement. We plot propagation time delay as a function of the coupling strength that can generate a phase shift of π radians across states.

**(B)** Unidirectional coupling model for nose twitching. Propagation time delay as a function of the coupling strength that can generate a phase shift of 0.65π radians across states.

### Supplementary Table Legends

#### Table S1. Electromyogram (EMG) recordings in each animal.

In all animals, thermocouple and head orientation sensor are implanted. L: Left muscle. R: Right muscle. SM: Sternomastoid. CM: Cleidomastoid. CT: Clavotrapezius. SP: Splenius. BC: Biventer Cervicis. VI: Vibrissa Intrinsic. NL: Nasolabialis. DN: Deflector nasi.

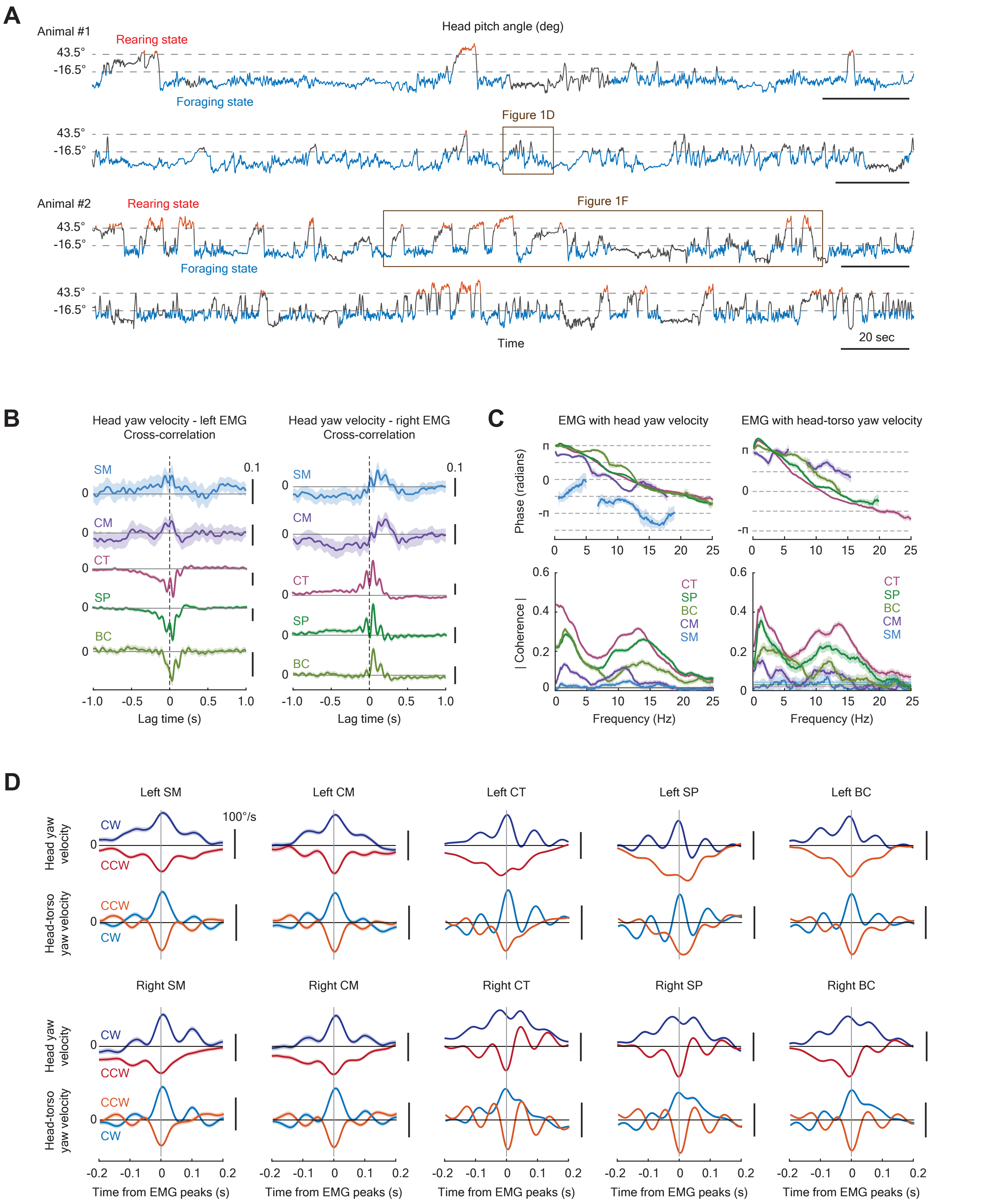


Figure S1. Liao and Kleinfeld (updated 15 Aug 2022)

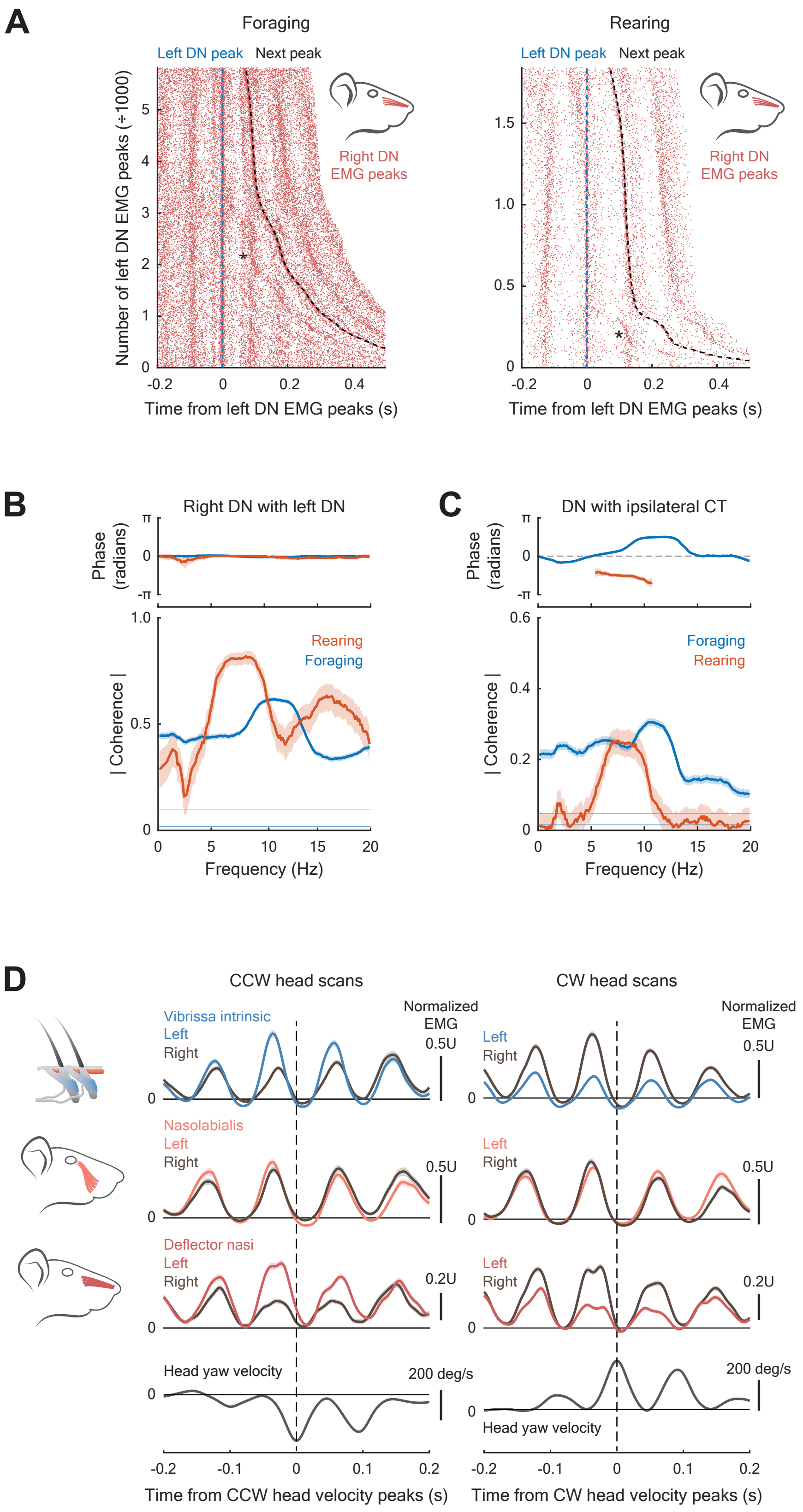


Figure S2. Liao and Kleinfeld (updated 23 November 2022)

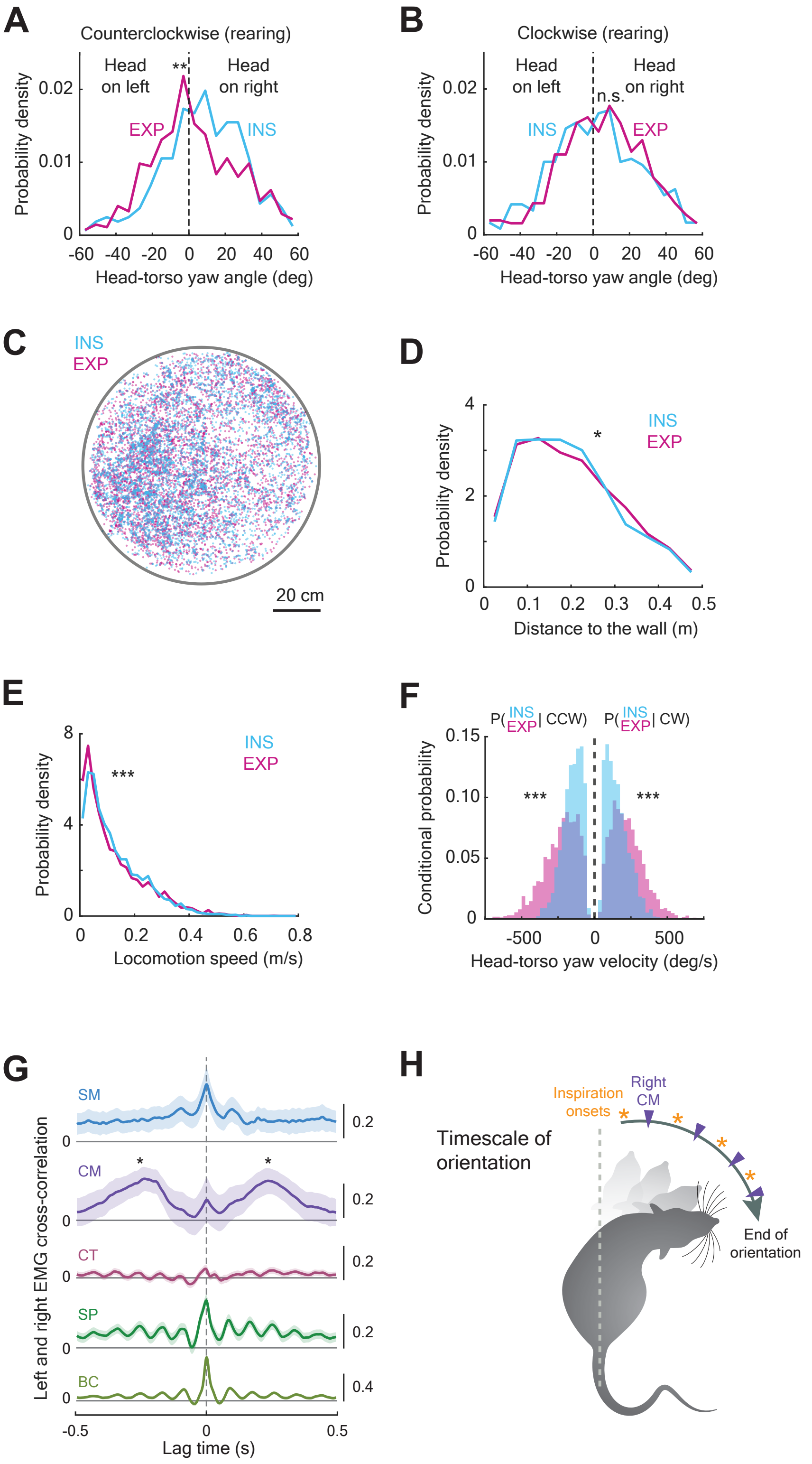


Figure S3. Liao and Kleinfeld (23 November 2022)

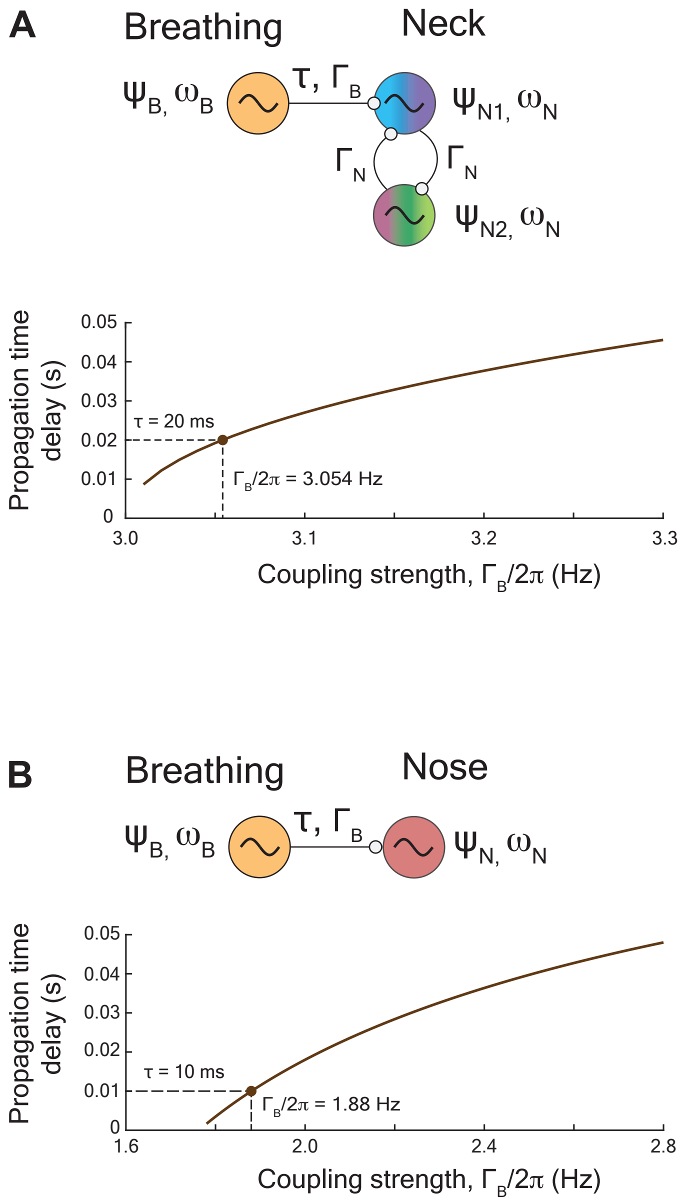


Figure S4. Liao and Kleinfeld (updated 18 December 2022)

**Table S1. Electromyogram (EMG) recordings in each animal.**

| # | ID | SM | CM | CT | SP | BC | VI | NL | DN | Torso sensor |
|---|---|---|---|---|---|---|---|---|---|---|
| 1 | SLR087 | | | | | | | | | X |
| 2 | SLR089 | | | | | | | | | X |
| 3 | SLR090 | | | | | | | | | X |
| 4 | SLR092 | | | | | | | | | X |
| 5 | SLR093 | | | | | | | | | X |
| 6 | SLR094 | | | | L + R | | | | | |
| 7 | SLR095 | | | | | | | | | X |
| 8 | SLR096 | | | L + R | L + R | | | | | |
| 9 | SLR097 | | | L + R | L + R | | | | | |
| 10 | SLR099 | | | | | | L | L + R | R | |
| 11 | SLR100 | | | | | | | | L + R | |
| 12 | SLR102 | | | L + R | | | | | | X |
| 13 | SLR103 | | | L + R | | | | | | X |
| 14 | SLR105 | | | L + R | | | | | | X |
| 15 | SLR106 | | | L + R | | | | | | X |
| 16 | SLR107 | | L + R | | | | | | | |
| 17 | SLR108 | | L + R | | | | | | | X |
| 18 | SLR110 | L + R | L + R | | | | | | | X |
| 19 | SLR111 | L + R | L + R | | | | | | | X |
| 20 | SLR112 | | | | L + R | L + R | | | | X |
| 21 | SLR113 | | | | L + R | L + R | | | | X |
| 22 | SLR114 | L + R | | | | | | | | |
| 23 | SLR115 | | | | L + R | L + R | | | | X |
| 24 | SLR116 | | | | | | L + R | | L + R | |
| 25 | SLR117 | | | | | | L + R | | L + R | |
| 26 | SLR119 | L | L | L | L | | | | | X |
| 27 | SLR120 | | | L | L | L | | | L | |
| 28 | SLR121 | | | L | L | L | | | L | |
| 29 | SLR122 | L | L | L | | | | | L | |
| 30 | SLR123 | L | L | L | | L | | | | |
| 31 | SLR124 | L | L | | L | L | | | | |
| 32 | SLR125 | | | | | | L + R | L + R | | |
| 33 | SLR126 | | | | | | L + R | L + R | | |

L: Left muscle. R: Right muscle. SM: Sternomastoid. CM: Cleidomastoid. CT: Clavotrapezius. SP: Splenius. BC: Biventer Cervicis. VI: Vibrissa Intrinsic. NL: Nasolabialis. DN: Deflector nasi.
